# Microclimatic heterogeneity is associated with forest structural complexity and biodiversity

**DOI:** 10.1101/2025.09.02.673839

**Authors:** Kerstin Pierick, Dominik Seidel, Pia Bradler, Simone Cesarz, Orsi Decker, Benjamin M. Delory, Sebastian Dittrich, Martin Ehbrecht, Nico Eisenhauer, Andreas Fichtner, Michael Junginger, Michael Köhler, Ludwig Lettenmaier, Roman M. Link, Soumen Mallick, Jörg Müller, Julia Rothacher, Lena Rugen, Guido F. Scholz, Jean-Léonard Stör, Kim K. Weißing, Clara Wild, Goddert von Oheimb, Christian Ammer, Bernhard Schuldt

## Abstract

Forest microclimates, their dependence on forest structure, and their impact on biodiversity are crucial for future forest management under climate change. However, standard approaches for measuring forest microclimates do not capture within-plot heterogeneity, which, according to the habitat heterogeneity hypothesis, is a key driver of local biodiversity.

We quantified horizontal and vertical microclimatic heterogeneity within 30 broad-leaved forest plots in Central Europe using a three-dimensional design with high spatial resolution. Moreover, we examined whether microclimatic heterogeneity differs among silvicultural treatments and whether it can be predicted using forest structure indices derived from laser scanning. Additionally, we explored the relationship between microclimatic heterogeneity and biodiversity.

In the understory of canopy gaps, warm and cold habitats co-existed in close proximity, leading to a high horizontal microclimatic heterogeneity. In closed stands with high structural complexity, we found steep gradients of increasing temperature and vapor pressure deficit from the ground to the canopy during mid-day. Canopy cover and forest structural complexity were strong indicators of microclimatic heterogeneity. We found positive relationships between herb layer temperature heterogeneity and the diversity of plants, Hymenoptera, and Diptera.

Our results demonstrate that single-point measurements fail to capture the substantial microclimatic heterogeneity within plots, potentially misrepresenting the conditions experienced by forest species. However, laser scanning provides reliable indicators for within- plot microclimatic heterogeneity. With canopy gaps featuring high horizontal microclimatic heterogeneity and promoting the biodiversity of several taxonomic groups, we argue that managing forests for maximized temperature buffering should not be the only strategy to conserve forest biodiversity.

**Highlights:**

- High small-scale horizontal microclimatic heterogeneity in canopy gaps
- Steep vertical microclimatic gradients in closed-canopy forests
- Canopy cover and structural complexity: indicators for microclimatic heterogeneity
- Positive relationship between herb layer temperature heterogeneity and biodiversity

## 1 Introduction

Forest microclimate has recently received a surge of interest, with a rapidly growing body of studies emphasizing its ecological importance, especially with regard to temperature buffering and the provision of cool microrefugia in the context of climate change (De Frenne et al., 2021, 2019; Haesen et al., 2023a, 2021). Compared to macroclimate, microclimate better represents the conditions that forest-dwelling organisms are experiencing (Haesen et al., 2023b; Kemppinen et al., 2024; Maclean and Early, 2023; Potter et al., 2013). Therefore, microclimate is central for understanding and predicting ecological patterns and processes from species niches and distributions (De Frenne et al., 2013; Gril et al., 2025; Lembrechts et al., 2019; Lenoir et al., 2017; Sanczuk et al., 2023), over community composition and diversity (De Frenne et al., 2013; Govaert et al., 2024; Seibold et al., 2016; Zellweger et al., 2020), to ecosystem functioning and forest dynamics (Beugnon et al., 2024; Braziunas et al., 2025; Hohnwald et al., 2020). However, the spatial resolution at which forest microclimate is usually measured is still relatively coarse compared to the scales relevant to forest-dwelling species (Pincebourde and Woods, 2020). In most studies, microclimate has been measured with one sensor per permanent plot at standardized heights, which was then assumed to be representative of the plot. Yet, depending on their structure, typical forest research plots (0.25 ha to 1 ha) likely provide a broader suite of microclimatic habitats than the one captured by a point measurement (De Frenne et al., 2025). To address this knowledge gap, here we investigate spatial microclimatic heterogeneity within forest plots.

Small-scale microclimatic heterogeneity, like other habitat heterogeneity in forests, can be separated into horizontal and vertical variability (Bae et al., 2025; Heidrich et al., 2020). Both have already been mapped before the advent of modern sensor and data logger technology (Geiger, 1927), but have, to our knowledge, barely ever been represented in the recent wave of microclimatic research. In fact, several recent studies have explicitly named vertical microclimate profiles as a major research gap (Bramer et al., 2018; De Frenne et al., 2021; Lenoir et al., 2017). In the few published examples where they were measured, steep vertical gradients in temperature and vapor pressure deficit were observed in forests (Link et al., 2020; Schuldt et al., 2011; Zahnd et al., 2023; Zweifel et al., 2002). Consequently, vertical microclimatic gradients provide a turnover in thermal habitats for canopy-dwelling arthropods (Brower et al., 2011; Saudreau et al., 2013). Furthermore, with their influence on the leaf gas exchange in different strata of tree crowns, vertical microclimatic gradients directly affect tree performance, and hence ecosystem-climate feedbacks (Bauerle et al., 2009; Vinod et al., 2023).

Not only vertical, but also horizontal microclimatic heterogeneity remains poorly understood, despite its potential importance for forest organisms. On the larger landscape scale, horizontal microclimatic heterogeneity varies between forest management systems (Menge et al., 2023). Small-scale heterogeneity of light availability, temperature, and soil moisture creates a mosaic of microhabitats on the forest floor, which increases understory plant diversity (Helbach et al., 2022), and the performance of ectotherms (Mueller et al., 2016; Woods et al., 2015). Previous works focused on describing small-scale horizontal or vertical microclimatic patterns (Ashcroft and Gollan, 2012; Gálhidy et al., 2006; Gray et al., 2002; Horváth et al., 2023; Zellweger et al., 2024). However, in order to compare microclimatic heterogeneity between forests and relate it to other variables, it is necessary to accurately quantify the microclimatic heterogeneity within observational units such as permanent plots.

The knowledge gap concerning spatial microclimatic heterogeneity is surprising, given how crucial heterogeneity is for biodiversity according to the habitat heterogeneity hypothesis. It states that heterogeneous habitats support the coexistence of more species with different niches (Eisenhauer et al., 2023; Heidrich et al., 2020; Huber et al., 2025; Stein et al., 2014). There is plenty of evidence for positive heterogeneity-diversity relationships (Heidrich et al., 2023; Helbach et al., 2022; Larrieu et al., 2015; Seibold et al., 2016). However, in most cases, heterogeneity of vegetation, land cover, or topography, but rarely (micro)climatic heterogeneity, have been investigated (Stein and Kreft, 2015). Landscape-scale structural heterogeneity is a key driver of biodiversity in Central European forests (Schall et al., 2018; Uhl et al., 2024), but effects can differ among taxonomic groups (Heidrich et al., 2020). Furthermore, the biodiversity of many taxonomic groups is differently affected by vertical and horizontal structural heterogeneity (Heidrich et al., 2020). The abundance of research on structural heterogeneity, compared to microclimatic heterogeneity, may stem from the fact that remote sensing enables efficient high-resolution measurement of forest structure, while spatially highly resolved, three-dimensional microclimatic measurements are still methodically challenging and resource-consuming (Zellweger et al., 2019b).

Forest structure is one of the key factors determining sub-canopy microclimate, together with macroclimate and topography (Borderieux et al., 2025; De Frenne et al., 2021; Zellweger et al., 2019a). Using remote sensing to model microclimate based on forest structure is therefore a promising application (Duffy et al., 2021; Frey et al., 2016; Gril et al., 2023; Kašpar et al., 2021; Vandewiele et al., 2023). Canopy cover, structural complexity, and vertical layering are key structural properties that directly impact ecologically relevant microclimatic variables such as air temperature, vapor pressure deficit (VPD), and soil moisture (Díaz-Calafat et al., 2023; Greiser et al., 2024, 2018; Kovács et al., 2017; Von Arx et al., 2013). If the relationships of these structural indices with microclimatic heterogeneity are as strong as with microclimate *per se*, inferring microclimatic heterogeneity from structural data would become straightforward (Zellweger et al., 2019b). This is a promising tool for modeling climatic microhabitats, and hence species distributions, biodiversity, and ecosystem functioning with even higher accuracy than based on microclimate *per se* (De Frenne et al., 2021; Lenoir et al., 2017).

Ultimately, the relationships between forest structure, microclimate, and biodiversity are central to the question of how biodiversity conservation should be integrated into forest management under climate change. Undoubtedly, forest management heavily influences forest microclimate via its impact on forest structures (Aussenac, 2000; Chen et al., 1999; Ehbrecht et al., 2019; Kovács et al., 2020). Silvicultural interventions leading to large canopy gaps, as well as natural disturbances that are currently increasing dramatically (International Tree Mortality Network, 2025), cause locally warmer conditions (Abd Latif and Blackburn, 2010; Gálhidy et al., 2006; Horváth et al., 2023; Thom et al., 2020). Structurally complex old-growth forests, on the other hand, have an especially pronounced buffering effect on sub-canopy microclimates (Frey et al., 2016; Wolf et al., 2021). While many recent studies on forest microclimate conclude that temperature buffering should be optimized by aiming at maximally dense canopies to maintain cool microhabitats under warming temperatures (e.g. De Lombaerde et al., 2022; Sanczuk et al., 2023; Thom et al., 2020), others argue that the special microclimate found in canopy gaps is important for enhancing biodiversity (Müller et al., 2023; Schall and Heinrichs, 2020). With this study, we aim to contribute to this debate by adding the aspect of small-scale microclimatic heterogeneity and its relationship with biodiversity.

We measured temperature, VPD, and volumetric soil water content (VWC) in an intensive three-dimensional design with 19 horizontally-distributed and nine vertically-installed loggers within each 50 m × 50 m plot, yielding 840 loggers in a total of 30 plots – covering a large vertical gradient from the soil to the canopy. We first describe the microclimatic patterns within plots. Next, we apply a statistical approach to summarize the vertical and horizontal small-scale heterogeneity of microclimatic variables per plot. We then test the hypotheses that (1) laser scanning-derived forest structure indices and microclimatic heterogeneity indices differ between experimental silvicultural treatments (Gap, Thinning, Control); (2) forest structure indices describing structural complexity, canopy cover, and vertical layering are good indicators for horizontal and vertical microclimatic heterogeneity; and (3) within-plot microclimatic heterogeneity is positively associated with the biodiversity of different taxonomic groups.

## 2 Material and methods

### 2.1 Study areas

The study was conducted at two sites in Germany that differ from each other in many aspects (Fig. 1a). The northern site, University Forest Sailershausen, is located in north-western Bavaria in the Haßberge region. At an elevation of 300 m – 360 m a.s.l., it is characterized by a mostly plane topography, and a comparatively warm and dry climate for Central European standards (during the reference period from 1981 to 2010, mean annual temperature 8.8°C and mean annual precipitation 732 mm (DWD Climate Data Center (CDC), 2025a, 2025b)). Formed on a complex mosaic of limestone bedrock with loess deposits, the soils are nutrient- and base-rich. The forest is owned by the University of Würzburg and managed for wood production according to the continuous cover forestry concept. Our plots are located in species-rich mixed broad-leaved forests dominated by ash (*Fraxinus excelsior*), European beech (*Fagus sylvatica*), hornbeam (*Carpinus betulus*), maple (*Acer* spec.), and oak (*Quercus* spec.) species. Maximum stand heights of the plots, as derived from mobile laser scanning (see section 2.4), were on average 25.9 m, with a standard deviation of 2.3 m.

**Figure 1:**
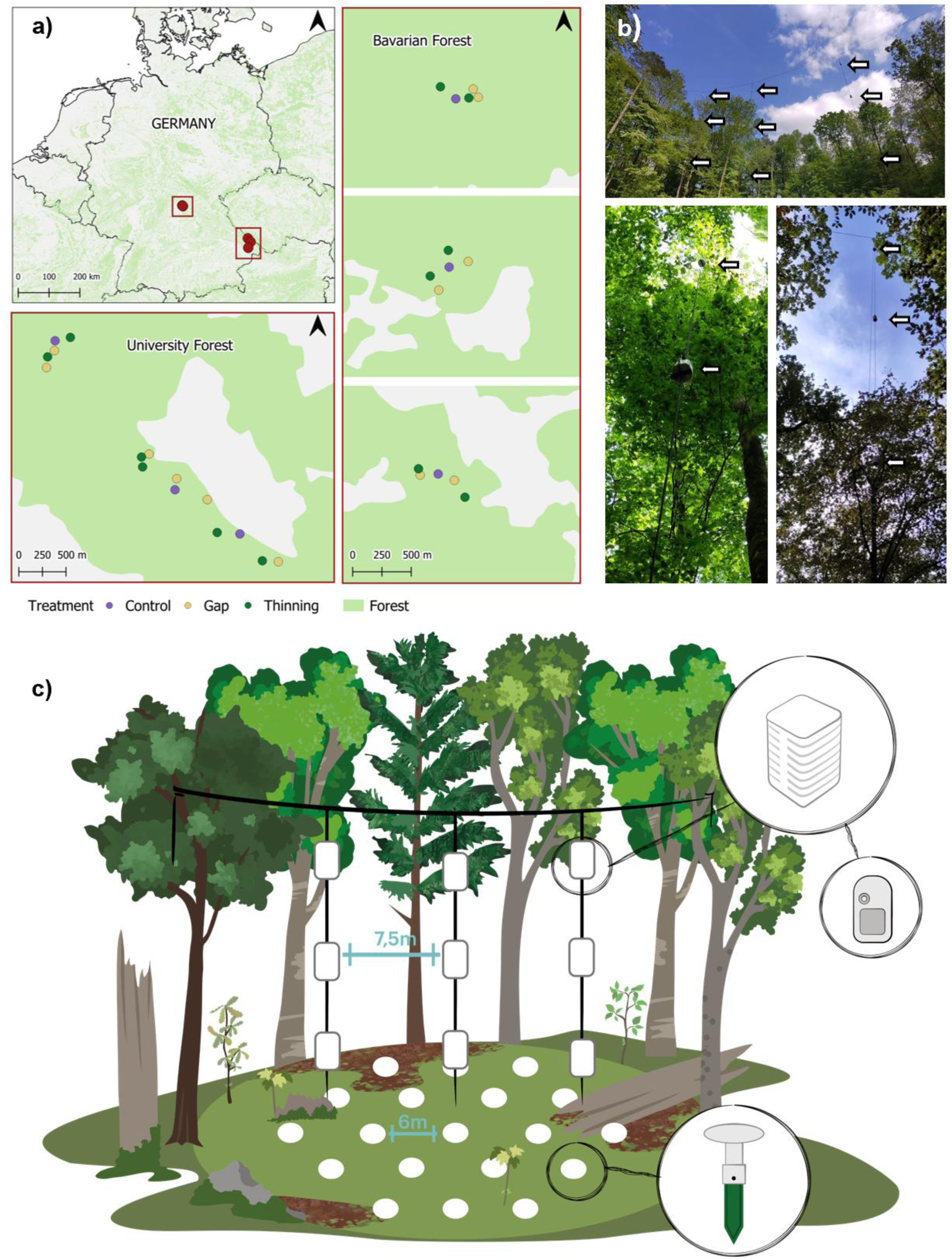
Experimental setup. a) Locations of the study sites, and spatial arrangement of the plots within the sites. b) Examples of loggers installed in sun shields on vertical ropes. Arrows highlight the positions of loggers. c) Logger installation scheme within the plots. The circles at the forest floor represent Tomst TMS-4 loggers, the rectangles along the ropes represent Onset Hobo MX2301A loggers installed within sun shields. Illustration by Rabea Klümpers.

The southern site is the Bavarian Forest low mountain range in south-eastern Bavaria. The study plots are clustered in three groups with elevations ranging from 490 m to 850 m a.s.l., and are mostly located on steep slopes. Accordingly, the climate covers a broad gradient but is generally colder and wetter than at the University Forest site (during to the reference period from 1981 to 2010, mean annual temperatures 5.9°C – 7.8°C and mean annual precipitation 1080 mm – 1416 mm, (DWD Climate Data Center (CDC), 2025a, 2025b)). Two plot groups are located within the management zone of the Bavarian Forest National Park, and the third is located outside the National Park, close to the city of Passau. All plot groups are located in forests that resemble typical broadleaf production forests of Central Europe. The soils, developed on gneiss and granite parent material, are more nutrient-poor and more acidic than at the other site. Our plots are located in beech forests with small shares of admixed Norway spruce (*Picea abies*). Maximum stand heights are 29.0 m on average, with a standard deviation of 2.3 m. At both sites, the stands were approximately 80-90 years old during our field campaigns.

### 2.2 Experimental design

We conducted the study in 30 plots with a size of 50 m × 50 m (0.25 ha), with 15 plots at the University Forest and 15 plots at the Bavarian Forest site. The plots are a subset of the plots established for the BETA-FOR experiment, a field experiment on the impact of forest structure manipulations on biodiversity and ecosystem functioning (Müller et al., 2023). In the course of experimental interventions in the winters of 2015/2016 (Bavarian Forest) and 2018/2019 (University Forest), ∼30% of the stand basal area was removed from treated plots, either aggregated in the center of the plot (“Gap”) or evenly distributed throughout the plot (“Thinning”), similar to silvicultural thinning operations. The canopy openings resulting from the Gap treatment had diameters of approximately 30 m. Both Gap and Thinning treatments were applied in two versions, either removing the entire trees (excluding the stumps) or retaining snags of 5 m height. However, since our key variables of interest were not significantly impacted by the presence of snags (Fig. S1), for this analysis, we pooled the treatment versions with and without snags. Additionally, Control plots without any interventions in the course of the experiment were established. This resulted in a total of six Control plots, and each twelve Gap and Thinning replicates, respectively (Fig. 1a).

### 2.3 Microclimatic measurements

The focus of this study lies on summer conditions, as this period is particularly relevant for microclimatic temperature buffering in forests (Steinparzer et al., 2025; Von Arx et al., 2012; Zellweger et al., 2019a) and represents a critical time span for many species in temperate forests, which are often in sensitive phases of their life cycles and exposed to heat stress during these months (Sallé et al., 2021; Teskey et al., 2015). Due to practical limitations, we could not measure microclimate at both sites in the same year. At the University Forest site, we measured from June 8 to September 30, 2023. In the following year, we measured during the same period (June 8 to September 30, 2024) at the Bavarian Forest site. During both measurement periods, average air temperatures at the respective sites were 1°C to 2°C warmer than the 30-year average from 1981-2010, with monthly precipitation sometimes exceeding, sometimes falling behind the long-term average ((DWD Climate Data Center (CDC), 2025a, 2025b, 2025c, 2025d), Fig. S2).

#### 2.3.1 Temperature and vapor pressure deficit in the tree layer

For measuring air temperature and relative humidity in higher stand layers, we installed three vertical ropes (either directly fixed to branches or to horizontal ropes) in the canopy of each plot (Fig. 1 b, c). One vertical rope was always positioned directly at the plot center, the other two 7.5 m north and south of the central rope, respectively. Depending on the local forest structure, we always aimed at installing the ropes as high as possible while still using branches stable enough to withstand the physical forces of the rope system. Along each rope, we installed three loggers, summing up to nine loggers per plot and a total of 270 devices. We could not use the data from six loggers at the University Forest site and eight loggers at the Bavarian Forest site due to technical malfunctions or storm damage on the trees holding the ropes. The lowest logger was installed at 5 m, the uppermost as high as the local conditions allowed, and the middle logger with approximately equal distances between the upper and lower ones. The maximum sensor heights per plot ranged from 12.5 m to 24.2 m. Logger heights were measured at the beginning and end of the measuring periods using a Vertex 5 ultrasound tree height meter (Haglöf, Långelse, Sweden). We used Onset Hobo MX2301A temperature and relative humidity loggers (Onset, Bourne, USA, from here on abbreviated as “Hobo loggers”) and protected them from direct solar radiation with TX Cover sun shields (Technoline, Bernburg, Germany). We measured temperature and relative humidity in 30-min intervals and calculated vapor pressure deficit (VPD) in kPa from these data as follows:

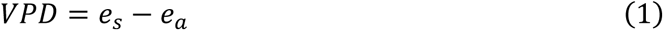

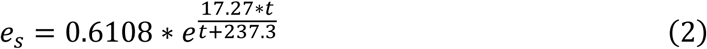

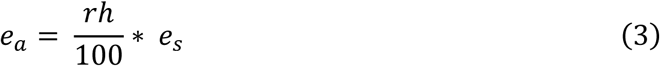

with *e_s_* being the saturation vapour pressure at a given temperature, *e_a_* the actual vapour pressure, *t* the air temperature in °C, and *rh* the relative air humidity. These measurements are referred to as “tree layer” measurements from here on.

#### 2.3.2 Temperature near the ground surface and soil moisture

To measure temperature near the ground surface and volumetric soil water content (VWC), we installed 19 standard Tomst TMS-4 devices (Wild et al., 2019) per plot. We arranged them in a hexagonal grid around the plot center, with the distance between each logger and its closest neighbors always being 6 m, and a maximal distance of 24 m for loggers at opposed corners of the grid (Fig. 1 c). Out of a total of 570 devices, four loggers from the University Forest (0.7%) and six loggers from the Bavarian Forest (1.1%) could not be used because they were missing or damaged after the measuring campaigns.

TMS-4 loggers (or “Tomst loggers” from here on) measure temperature -6 cm beneath the soil surface, 2 cm above the soil surface, and 15 cm above the soil surface, as well as a proxy for VWC in the uppermost horizon (Wild et al., 2019). The measurement interval was set to 15 min. We refer to the measurements at 15 cm height as “herb layer” measurements. For processing the data, we used the R package myClim 1.4.0 (Man et al., 2023). VWC was derived from the raw logger output using calibration parameters based on soil texture (see Method S1 for details).

### 2.4 Measuring forest structure

We used mobile laser scanning (MLS) to capture the three-dimensional forest structure of the plots and to derive forest structural indices as potential indicators of microclimatic heterogeneity. Each plot was scanned in the same summer (July-August) as the microclimatic measurements using a ZEB Horizon laser scanner (GeoSLAM Ltd., Nottingham, UK). Starting at the plot center, we carried the handheld laser scanner through the plot in outward spirals, circling the plot center with an increasingly larger distance until reaching 30 m, then returning to the center to close the scan loop. The resulting point clouds were processed in the Faro Connect software (Faro Technologies Inc., Lake Marry, USA) to perform the simultaneous locating and mapping (SLAM) using the “forest” environment as presetting. The resulting .laz- files were subsampled to 1 cm resolution, noise filtered to remove outliers, subsequently normalized to eliminate terrain effects and finally exported as xyz-files. Based on these, we calculated three forest structural indices: Box dimension, canopy cover, and effective number of layers (ENL).

The first, box dimension, is a measure for forest structural complexity based on the fractal dimension of the stand. It is a dimensionless number ranking the structural complexity between 0 (no complexity) and 3 (a solid filled cube of plant material), with both the amount of material and its distribution in space affecting the final value (Basnet et al., 2025; Mandelbrot, 1977; Neudam et al., 2022; Seidel, 2018).

The second index, canopy cover, expresses the fraction of the area vertically covered by vegetation elements (at least one laser hit), calculated based on a 20 cm × 20 cm ground resolution and considering all points higher than 5 m above ground as “canopy” (e.g., Höwler et al., 2024). Canopy cover is given in percent and serves as a proxy for the availability of direct light to the lower forest strata.

Finally, ENL was calculated as introduced by Ehbrecht et al. (2016) for laser scanning data. This measure describes the vertical layering of the forest via the occupation of 1 m layers with points, analogously to describing diversity with Hill numbers (Hill, 1973). Here, we calculated ENL1D, i.e., weighting the layers by their occupancy as in Hill numbers of q = 1 (Ehbrecht et al., 2016; Hill, 1973).

### 2.5 Biodiversity

Using data from the BETA-FOR experiment (Müller et al., 2023), we explored how different taxonomic groups respond to small-scale microclimatic heterogeneity. From a larger set of available biodiversity data, we selected four exemplary clades spanning different levels of mobility: vascular understory plants, spiders, Diptera (flies), and Hymenoptera (sawflies, bees, wasps, and ants). Importantly, we selected them before inspecting the data and did not look into more taxonomic groups than the ones mentioned here.

Vascular plant species were recorded in five relevés of each 50 m² per plot during the vegetation period of 2023. We recorded the incidence of all herbaceous and woody species in the understory up to a height of 1 m.

Spiders were sampled with two pitfall traps per plot installed at a distance of 7 m from the plot center, one in the northwest and the other in the southeast, from May to July 2022. The trap consisted of a 400 ml plastic cup buried at ground level. Pitfall traps were covered with transparent plastic roofs (15 × 15 cm, approximately 5 cm above the ground) to protect the trap from rain or leaves. As trapping liquid, we used a ∼15 % solution of NaCl and a drop of odorless detergent to break the surface tension, which ensured a rapid killing of sampled arthropods. The traps were emptied monthly, sorted to order level, and spiders were identified by an experienced taxonomist. We aggregated the data to the plot level by summing up the abundances per species from all 180 samples (i.e., 30 plots × 2 traps × 3 months).

Diptera and Hymenoptera were sampled using one Malaise trap per plot. Traps operated from the beginning of May until the end of July 2022 and were equipped with collection bottles filled with ethanol (70%). Samples were collected fortnightly, separated into two size classes and sequenced for taxonomic identification on the OTU level (for details on the metabarcoding process, see Rothacher et al. (2025)). We filtered the OTUs for Diptera and Hymenoptera, and aggregated the samples to the plot level by summing up the reads for each OTU.

### 2.6 Data analysis

#### 2.6.1 Continuous predictions over sensor heights

In order to describe temperature and VPD at different heights during the day, we fitted four generalized additive models (GAM), one for the two microclimatic variables air temperature and VPD for each of the two study sites, respectively. We fitted separate models for the two sites because not only do they differ in climate, but also they were measured in different years. The GAMs were fitted using the R package `mgcv` version 1.9.1 (Wood, 2011). We modeled the climatic variable in dependence of treatment, plot, plot-specific thin plate regression splines along the continuous time during the measurement period using k = 25 knots, and treatment-specific tensor product smooths containing a cubic regression spline basis for sensor height, a second cubic regression spline basis for continuous time during the measurement period, and a cyclic cubic regression spline basis for time throughout the day. Accordingly, we assumed the microclimatic variables to differ on average between treatments and plots, with a plot-specific long-term trend and a treatment-specific diurnal pattern that was permitted to change over the course of the measurement period and to differ depending on sensor height. We then generated predictions for each combination of the variables ‘day of the measurement period’, ‘treatment’, ‘time of the day’, and ‘sensor height’. To acquire predictions for an average summer day, we calculated the average prediction for each time of the day over all days during the study period. These models were only used for graphically summarizing the highly resolved data, not for statistical hypothesis testing.

#### 2.6.2 Quantifying microclimatic heterogeneity

To quantify horizontal within-plot heterogeneity of the microclimatic variables measured with Tomst loggers (temperatures at -6 cm, 2 cm, 15 cm, and topsoil VWC), we calculated the standard deviation of the 19 measured values per plot and variable at each point in time (i.e., every 15 min over the study period of almost four months). Then, we calculated the plot averages of these standard deviations over the whole study period.

For the Hobo loggers installed on ropes, we wanted to disentangle horizontal and vertical heterogeneity. We first aggregated the measured time series to the logger level by calculating the average daily maximum temperature and VPD of each Hobo logger. We focused on daily maximum values because they are crucial for organisms in terms of heat and drought stress (Bristow and Abrecht, 1991; Macek et al., 2019). Additionally, we repeated the analysis with average daily mean instead of maximum values and obtained similar results, since average daily mean and maximum temperatures were highly correlated (results not shown). As a next step, we fitted one linear model per plot, where the nine average daily maximum values per plot were modeled in dependency of height (in m) and horizontal position (a factor with three levels indicating on which of the three ropes the logger was installed). We then performed variance decompositions of each of the models using the Lindeman, Merenda, and Gold (LMG) approach to variance decomposition (Lindeman et al., 1980). By doing so, we obtained the variance explained by the linear vertical gradient, the variance explained by the horizontal position, and the unexplained variance (differences between logger readings not explained by either horizontal or vertical position). We interpret the explained variance components as horizontal and vertical heterogeneity of average daily maximum temperature and VPD in the tree layer. Since the three components (horizontal, vertical, and unexplained) add up to 100%, our horizontal and vertical heterogeneity indices are not entirely independent, but can vary freely by some degree through the unexplained component.

#### 2.6.3 Relationships between treatment, forest structure, and microclimatic heterogeneity

We tested whether the forest structure indices (box dimension, canopy cover, ENL) and the microclimatic heterogeneity variables differed between the Gap, Thinning, and Control treatments using Kruskal-Wallis tests and post-hoc Dunn tests. These non-parametric tests were chosen because variance homogeneity within groups was not given in most cases. For the Dunn tests, we used the R package `dunn.test` version 1.3.6 (Dinno, 2024). Furthermore, we tested whether the microclimatic heterogeneity variables depended on forest structure indices using linear models. We fitted one separate model for each pairwise combination of response and predictor and did not include additional covariates such as site or treatment, since we were interested in the usefulness of the forest structure indices as proxies for microclimatic heterogeneity without any additional information.

#### 2.6.4 Relationships between microclimatic heterogeneity and biodiversity

We used rarefaction curves to estimate the sample coverage of the measured data and to standardize the taxonomic species richness data to identical sample coverages by inter- or extrapolation. This was computed using the `estimate3D` function from the `iNEXT.3D` R package version 1.0.6 (Chao et al., 2021), run in R version 4.3.0 (R Core Team, 2025). The taxonomic species richness data were standardized to sample coverage values of 0.90 for vascular plants, 0.95 for spiders, 0.996 for Diptera, and 0.997 for Hymenoptera. These values were chosen to acquire a majority of the data from interpolation, not extrapolation. The differences result from varying estimated sample coverages of the measured data between the taxonomic groups. For more information, see Rothacher et al. (2025).

Since the biodiversity data were strongly dominated by differences between the two sites, with all taxa being more diverse at the University Forest site (Fig. S3), we first fit a linear model with site as predictor for each taxonomic group and extracted the residuals. We then used the scaled residuals as site-corrected biodiversity data and fitted linear models for each pairwise combination of microclimatic heterogeneity variable and taxonomic group. If not stated otherwise, the data analysis was conducted in R version 4.5.0 (R Core Team, 2025).

## 3 Results

### 3.1 Within-plot microclimatic patterns

Air temperatures showed typical diurnal oscillations at all heights, with maximum temperatures around 15:00 CET during the observation period from June to September. During this time of the day, differentiation of temperatures along vertical profiles and between treatments was most pronounced (Fig. 2 a, b, c). During the evening, night, and morning, air temperatures in the tree layer (> 5 m) on average showed no clear vertical gradients, but during the afternoon, they increased from lower to higher air layers in thinned and control plots (Fig. 2 a, Fig. S4). At the University Forest site, average temperatures in the Control treatment at 15:00 increased from 22.5°C at 5 m to 24.1°C at 20 m (Fig. 2 a). In the canopy gaps, no such pronounced vertical gradient existed: there, the average predicted temperatures at 5 m height were 23.7°C, and at 20 m height 24.3°C. As a result, air temperatures in the Gap treatment were warmer than in the Control, especially during the noon and afternoon hours, and only in the lower air layers, with differences between 1°C and 2°C (Fig. 2 b). At the Bavarian Forest site, apart from overall lower air temperatures, the patterns were similar, but vertical temperature gradients in the Control and Thinning treatments were less pronounced, and the gaps did not differ as much from Control during the early afternoon. Vapor pressure deficit (VPD) displayed similar patterns as temperature, with the exception that a vertical gradient was also present in the gaps (but less pronounced than in the thinned and control plots, Fig. S4). Individual vertical profiles of temperature and VPD were variable and partly differed from the predicted average patterns (Fig. 2c, S5).

**Figure 2:**
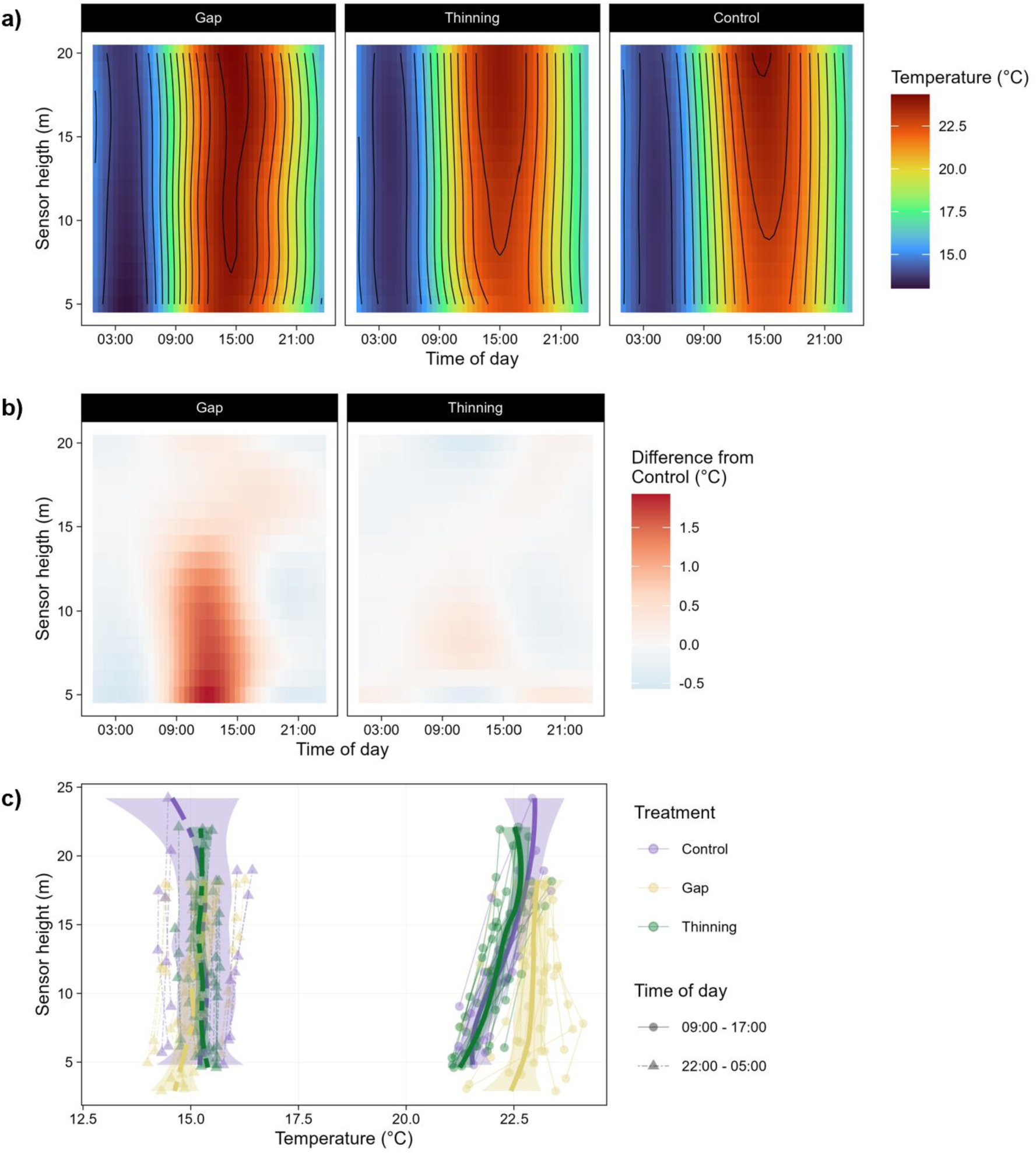
Temperature patterns in the tree layer (shown are only data for the University Forest due to the large climate differences between the two sites that preclude pooling). a) Estimated average air temperatures in the tree layer per height, time of the day, and treatment. Shown are predictions from a generalized additive model (GAM). b) Temperature differences between Control and the other treatments (Gap and Thinning) for different heights throughout the day. We subtracted the GAM predictions shown in Fig. 2a) for the control plots from the predictions of each of the two treatments to illustrate the magnitude of the differences. c) Average day and night vertical profiles. Each point is one logger, with the thin lines connecting loggers installed on the same rope. The bold lines are loess smoothers for each group along the y axis. For equivalent results from the Bavarian Forest site and vapor pressure deficit (VPD), please see Fig. S4 and S5.

Temperatures close to the ground surface also showed the clearest vertical stratification and differentiation between treatments in the afternoon (Fig. 3 a). During these hours, average temperatures increased from Control over Thinning to the Gap treatment at all measuring heights and at both sites, even though the differences were always smaller than 1°C (Fig. 3 a, Fig. S6 a). The temperatures measured simultaneously within the same plot and at the same height could vary by several degrees (in °C) in some examples, but were relatively homogeneous in others. For example, in plot 2, a gap, average temperatures at 15:00 ranged from 21.8°C to 27.5°C, whereas in plot 60, a thinned plot, they only ranged from 21.5°C to 22.2°C (Fig. 3 b). Volumetric water content of the topsoil (VWC) did not oscillate diurnally like temperatures and VPD, but suddenly increased during precipitation events and then slowly decreased until the next rainfall. In the Bavarian Forest, Control displayed lower VWC than the other two treatments in the early measurement period. In the University Forest, the gaps were drier than the closed stands in the early phase, but wetter towards the end of summer (Fig. S6 b).

**Figure 3:**
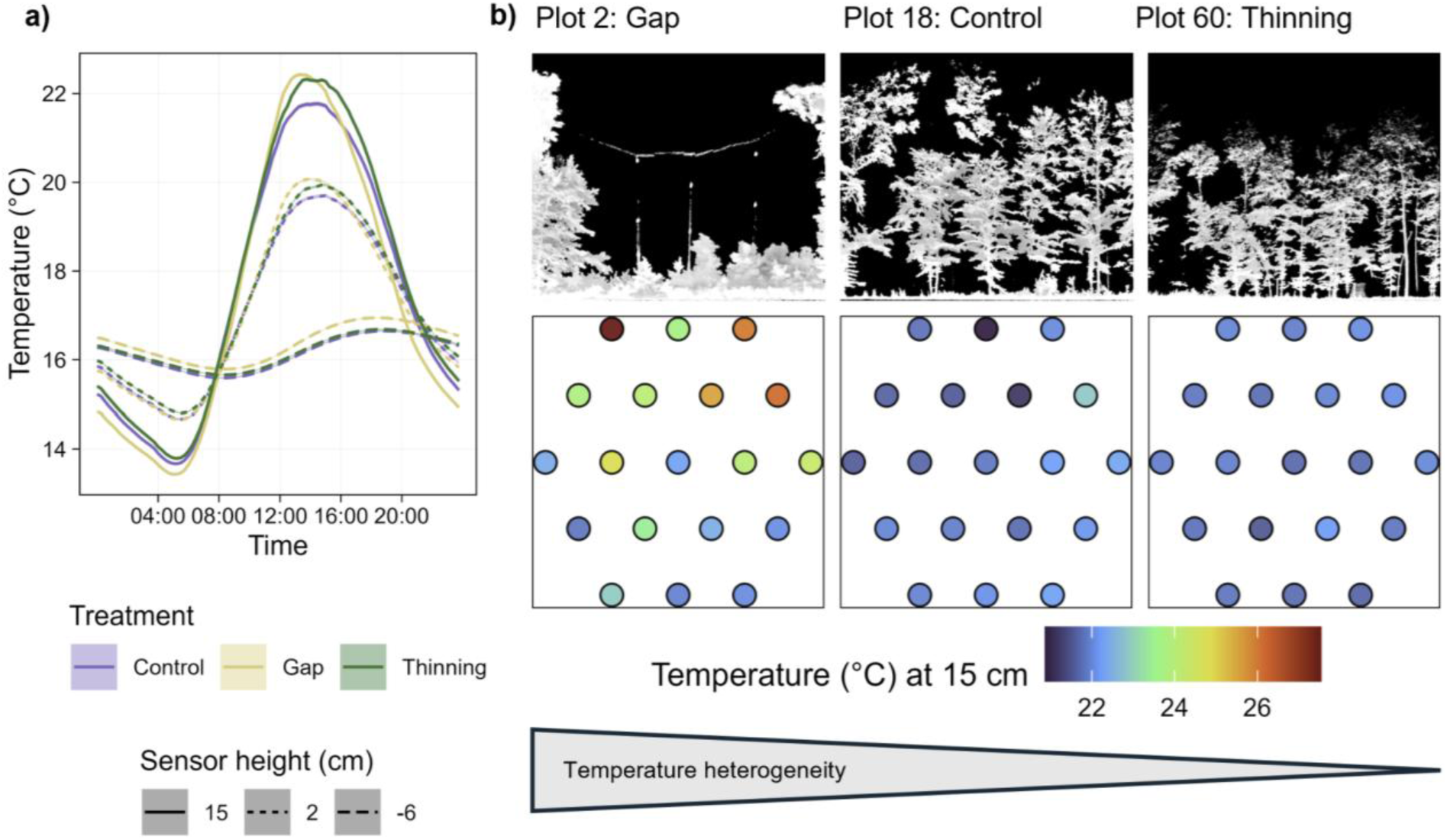
Temperature patterns in the herb layer. a) Mean temperatures close to the soil surface, measured with 19 Tomst loggers per plot, averaged over all loggers per treatment and all days of the observation period. b) Average temperatures in the herb layer (15 cm height) at 15:00, the approximate time of daily maximum temperatures, in three selected exemplary plots. Each point represents one Tomst logger, with their positions in the plot being shown from a bird’s perspective. In the top row, projections of cross sections through the 3D point clouds illustrate the forest structure of the selected plots. The three exemplary plots were selected to represent high, intermediate, and low horizontal temperature heterogeneity. For results from both sites, please see Fig. S6.

### 3.2 Effects of the treatments on forest structure and microclimatic heterogeneity

Box dimension and canopy cover, but not effective number of layers (ENL), differed significantly between the experimental treatments. In gaps, box dimension and canopy cover were significantly lower than in the other treatments, while they were not significantly different between thinned and control plots (Fig. 4 a, Fig. S7, Tab. S1). Canopy cover and box dimension were highly correlated with each other (*r* = 0.88), while ENL was independent from them (Fig. S7).

**Figure 4:**
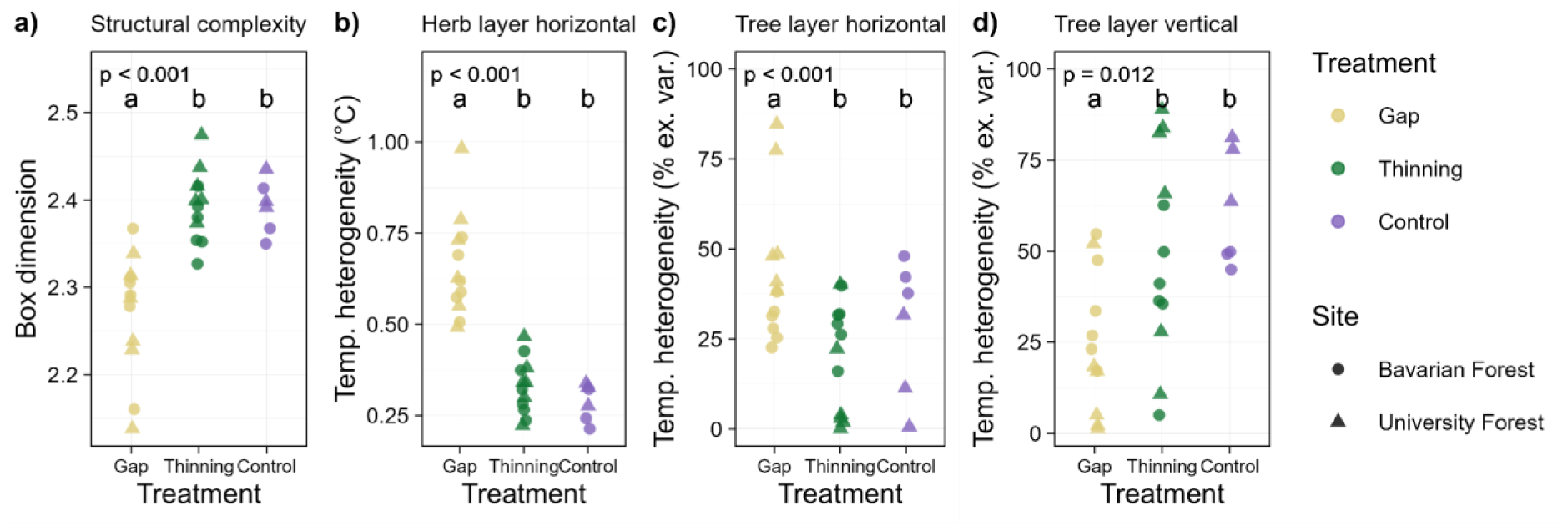
Effect of treatments on forest structural complexity and within-plot temperature heterogeneity. Letters indicate whether groups significantly differ from each other. This was tested with Kruskal-Wallis tests and post-hoc Dunn’s tests, see Tab. S1 for the full test output. The p-values are from the Kruskal-Wallis tests, while the letters indicate which groups differed from each other according to the Dunn’s tests. The abbreviated y axis labels stand for temperature heterogeneity, either quantified as average within-plot standard deviation (°C) or as % explained variance (% ex. var.). See Fig. S7 and S8 for results for all forest structure and microclimatic heterogeneity variables.

Horizontal temperature heterogeneity close to the soil surface (-6 cm, 2 cm and 15 cm) was higher in the Gap treatment than in Thinning and Control, but treatment had no effect on VWC heterogeneity (Fig 4 b, Fig. S8, Tab. S1). Vertical heterogeneity of temperatures and VPD in the tree layer were significantly lower in the gaps than in thinned and control plots, whereas horizontal temperature heterogeneity in the tree layer was significantly higher in gaps than in thinned plots. For none of these variables, the difference between Thinning and Control was significant (Fig. 4 c, d, Tab. S1). Treatment had no effect on horizontal VPD heterogeneity (Fig. S8, Tab. S1).

### 3.3 Effects of forest structure on microclimatic heterogeneity

Box dimension and canopy cover were significantly associated with all microclimatic heterogeneity variables except VWC heterogeneity, which had no significant relationship with any predictor (Fig. 5, Fig. S9, Tab. S2). ENL did not have significant effects on any of the response variables. The relationships of box dimension and canopy cover with all variables describing horizontal heterogeneity were negative, i.e., the higher the structural complexity and denser the canopy, the more horizontally homogeneous were the temperatures and VPD. In contrast, the relationships of the vertical heterogeneity variables with box dimension and canopy cover were positive, i.e., in structurally more complex plots with denser canopies, vertical gradients were steeper (Fig. 5, Fig. S9, Tab. S2). Of the significant relationships, in soil temperature heterogeneity (-6 cm) the least amount of variance was explained by the predictors (16% for box dimension, 17% for canopy cover). The highest fraction of variance levels was explained for horizontal herb layer temperature heterogeneity (70% for box dimension, 79% for canopy cover). All other models with significant relationships explained between 28% and 49% of the variance in the data, with canopy cover usually explaining slightly more variance than box dimension (Tab. S2).

**Figure 5:**
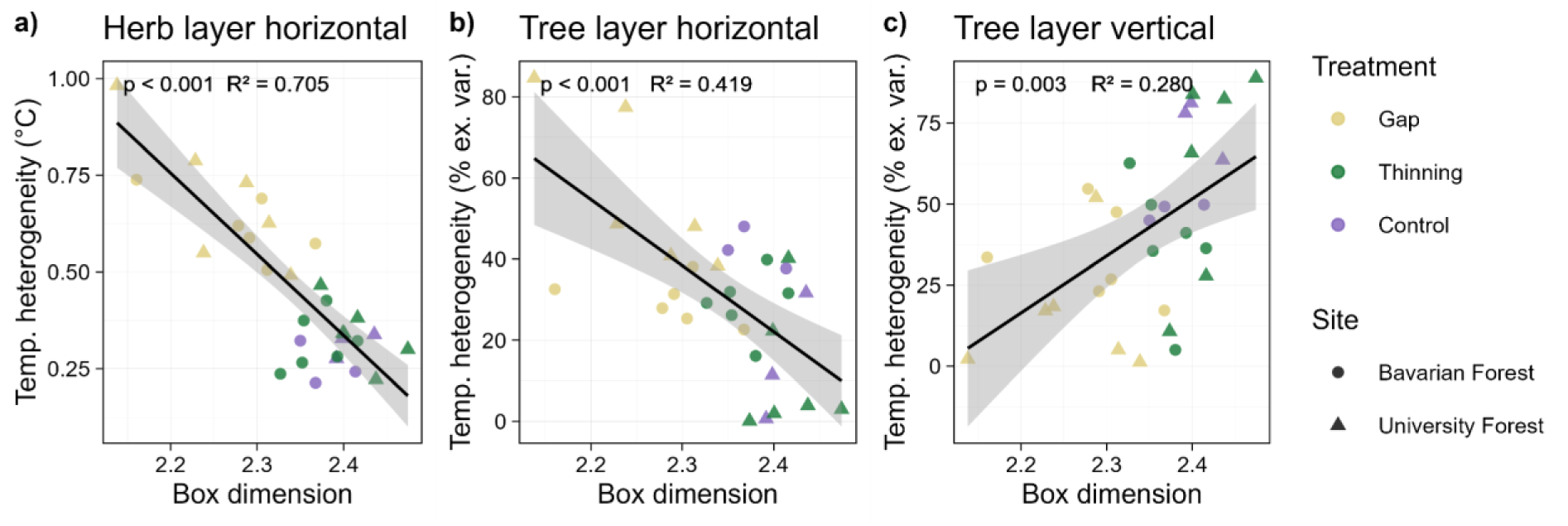
Temperature heterogeneity indices in dependency of forest structural complexity (quantified as box dimension). The lines represent predictions from linear models with 95 % confidence intervals. The slope parameters were significantly different from 0 for all three pairwise relationships. See Tab. S2 for model results. Given are the p-value of the slope and the R^2^ value of the linear regression. The abbreviated y-axis labels stand for temperature heterogeneity, either quantified as average within-plot standard deviation (°C) or as % explained variance (% ex. var.). See Fig. S9 for results for all forest structure and microclimatic heterogeneity variables.

### 3.4 Relationships between biodiversity and microclimatic heterogeneity

While heterogeneity of microclimatic conditions in higher strata and volumetric soil water content (VWC) heterogeneity were not significantly associated with biodiversity, heterogeneity of temperatures near the soil surface was significantly associated with increased diversity of vascular plants, Diptera, and Hymenoptera (Fig. 6, Fig. S10, Tab. S3). The relationships were significant for temperatures at 15 cm in all three groups, at 2 cm for Hymenoptera and vascular plants, and at -6 cm just for vascular plants. The models explained between 14% and 39% of the variance in the data. Spider diversity was not significantly associated with any of the variables (Tab. S3).

**Figure 6:**
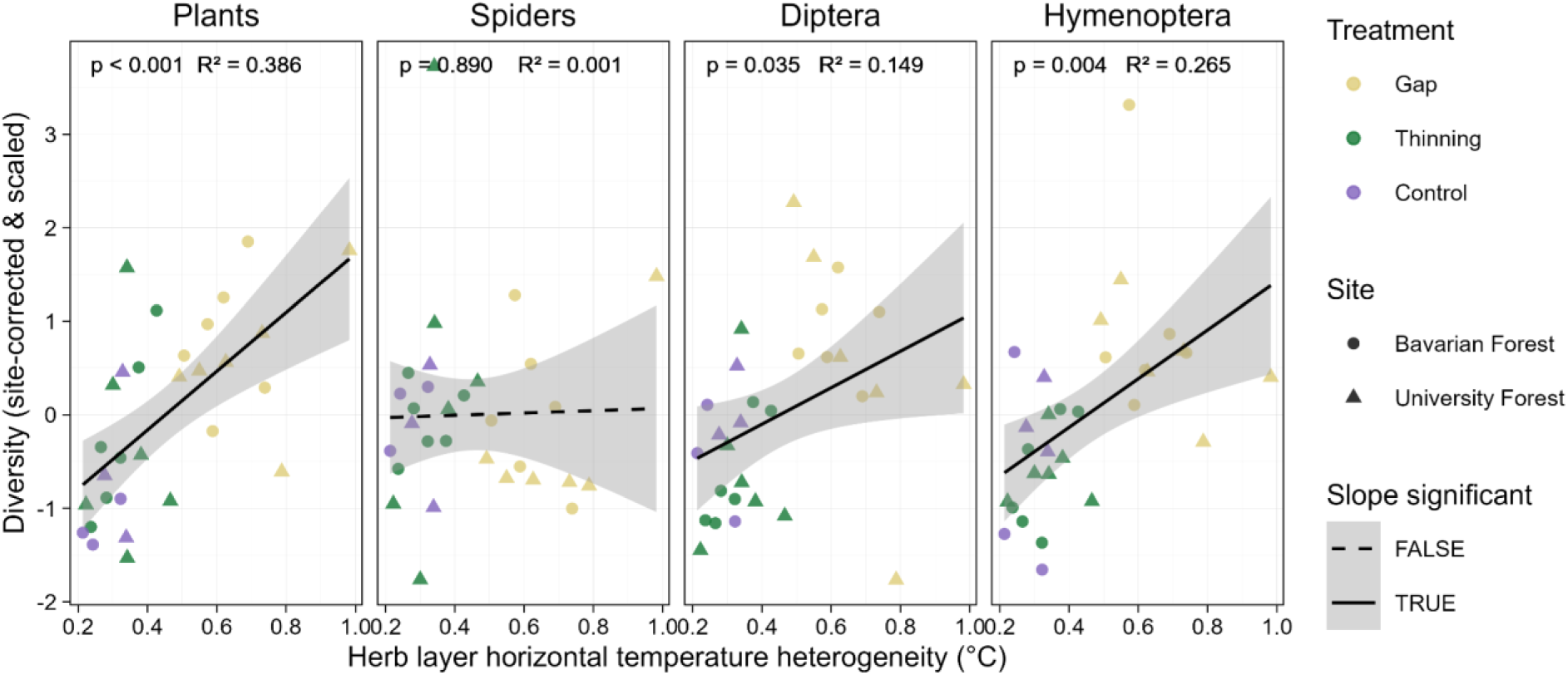
Relationships between within-plot temperature heterogeneity at 15 cm height and diversity of selected taxa. Shown are the data together with predictions and 95 % confidence intervals from linear models, as well as the p-value of the slope and the R^2^ value of the linear regression. See Tab. S3 for model results and Fig. S10 for relationships between other microclimatic heterogeneity variables and biodiversity.

## 4 Discussion

By installing an intensive design of sensors in the three-dimensional space in forest plots, we could quantify horizontal and vertical microclimatic heterogeneity within plots, compare them between silvicultural treatments, and test whether they can be predicted with laser scanning-derived forest structure indices. We then explored whether horizontal and vertical microclimatic heterogeneity are associated with the biodiversity of various taxonomic groups.

### 4.1 Microclimatic heterogeneity in gaps and closed stands

Our first hypothesis, stating that microclimatic heterogeneity varies between canopy gaps, thinned stands, and control plots, was partly supported by our results: gaps differed both in terms of horizontal and vertical microclimatic variability from the two other treatments. However, there was no significant difference between thinned and control plots. Forest structural complexity and canopy cover showed the same pattern. A high degree of structural similarity between thinned and control plots in this experiment was also found using other remote sensing platforms and forest structure indices, and is presumably caused by crown expansion of the remaining mature trees after the thinning operations (Kacic et al., 2025, 2024; Pierick et al., 2025). Therefore, differences in microclimatic heterogeneity that might have been present directly after the interventions did not prevail.

We found steep vertical gradients of air temperature and vapor pressure deficit (VPD) during midday in closed-canopy stands, with a gradual decrease from warmer and drier air in the upper strata of the forest to cool, moist conditions towards the forest floor. It is well established that this is a result of large proportions of the incoming short-wave radiation being absorbed or reflected by the upper strata of the tree crowns (Geiger, 1927). In gaps, the solar radiation penetrates to the understory vegetation or forest floor, where the energy transfer from short- to longwave radiation takes place. The emitted long-wave radiation in combination with air mixing leads to comparatively warm, vertically homogeneous air temperatures in the tree layer (Geiger, 1927). Previous measurements (though with fewer replicates) yielded similar results concerning vertical microclimatic profiles in forests (De Frenne et al., 2021; Geiger, 1927; Link et al., 2020; Schuldt et al., 2011).

As expected, the horizontal heterogeneity of temperature and VPD was higher in gaps than in closed stands, especially temperature heterogeneity in the herb layer. This was driven by closed stands being homogeneously cold during the diurnal maximum temperatures, while gaps featured cooler and warmer temperatures in close proximity. It is trivial and has long been known that, during the vegetation period, midday air temperatures and VPD are on average higher in canopy gaps than in closed stands (Abd Latif and Blackburn, 2010; Chen et al., 1999; Geiger, 1927), but the coexistence of colder and warmer microhabitats within gaps has typically been overlooked (but see Gálhidy et al. (2006), Gray et al. (2002), and Horváth et al. (2023) for intricate maps of microclimatic mosaics within canopy gaps). The two most likely contributing factors are north-south gradients of shading by the gap edges (Gray et al., 2002; Horváth et al., 2023; Ritter et al., 2005), and, to a greater degree, understory vegetation emerging after the opening of the canopy. Our measurement campaigns took place five and nine years after the interventions, leaving enough time for herbs, shrubs, and regenerating trees to form a dense understory as a reaction to the increased light availability (Pierick et al., 2025; Thom et al., 2022; Zhu et al., 2014). This provides patches of deep shade even under otherwise hot gap conditions (Brůna et al., 2024; Stickley and Fraterrigo, 2021). Sun flecks, while being crucial for the energy balance at the forest floor and briefly causing extreme small-scale differences (Chazdon and Pearcy, 1991; Way and Pearcy, 2012), could not provide the same degree of thermal heterogeneity under closed canopies (but see Mueller et al. (2016)).

We found no impact of the treatments on VWC heterogeneity. Previous studies reported effects of canopy openings on soil moisture (Belmonte et al., 2022; Horváth et al., 2023), driven by opposing mechanisms of increased evaporation from the soil, but decreased interception and transpiration by vegetation (Abd Latif and Blackburn, 2010). Precise measurements of soil moisture are challenging and can be biased depending on the sensors used (Jackisch et al., 2020). The signal from Tomst TMS-4 loggers used in this study needs to be calibrated based on soil texture information (Wild et al., 2019). Since we assumed soil texture to be homogeneous within our plots, our calibrated data might underestimate true VWC heterogeneity. Furthermore, the influence of other unmeasured factors, such as topography, might have masked potential signals of forest structure (Chaney et al., 2015).

### 4.2 Forest structure indices as proxies for microclimatic heterogeneity

Our second hypothesis was that forest structure indices quantifying forest structural complexity, canopy cover, and vertical layering can be used as indicators for within-plot microclimatic heterogeneity. Our results show that especially temperature heterogeneity in the herb layer is strongly associated with forest structural complexity (as quantified by the box dimension index) and canopy cover. Even though we pooled data from two dissimilar forests that were measured in different years, bivariate linear models of herb layer temperature heterogeneity in dependency of box dimension or canopy cover explained over 70 % of the variance. While canopy cover explained more variance than structural complexity for most variables, the differences were minor, since canopy cover and box dimension were highly correlated. We still consider structural complexity the more useful indicator for microclimatic heterogeneity, because it distinguishes better between dense stands. Even under canopies with > 95 % cover, space filling—and thus structural complexity—may vary greatly. These structural differences, being potentially important for sub-canopy microclimates, are captured by the box dimension. The poor performance of vertical layering (ENL) as a proxy for microclimatic heterogeneity in our dataset likely results from the pronounced differences between dense stands and large canopy gaps that characterizes our study design. In other datasets, where only dense stands are compared with each other, the vertical layering might be more relevant for below-canopy microclimates (Ehbrecht et al., 2019; Gril et al., 2023). Therefore, box dimension, integrating both the vertical and horizontal distribution of vegetation in space, is in our view the most promising proxy for local microclimatic heterogeneity, with good chances of it being universally applicable to datasets spanning both broad (as in this study) and narrow gradients of canopy cover. This remains to be tested in future studies.

While we used mobile laser scanning (MLS), i.e., ground-based measurements with a hand- held device, to capture the three-dimensional forest structure, other remote sensing methods will likely provide similarly useful predictors for microclimatic heterogeneity. Indices derived from airborne laser scanning that quantify similar structural characteristics as our MLS-derived indices, e.g., canopy openness and the vertical complexity index, are useful predictors for forest microclimate (Gril et al., 2023; Menge et al., 2023; Vandewiele et al., 2023). Even metrics from spaceborne remote sensing are highly correlated with the MLS-derived indices used in this study (Kacic et al., 2025). This suggests that such forest structure metrics can be used to infer vertical profiles and small-scale horizontal microclimatic heterogeneity, opening the possibility to map microclimatic habitats in landscapes with even greater precision (Zellweger et al., 2019b).

### 4.3 Associations of microclimatic heterogeneity with biodiversity

In line with our third hypothesis, we found positive associations between horizontal temperature heterogeneity in the herb layer and the biodiversity of vascular plants, Diptera, and Hymenoptera. One possible explanation for this pattern is that, as predicted by the habitat heterogeneity hypothesis, forest plots with more heterogeneous thermal conditions could harbor more species with contrasting niches (Eisenhauer et al., 2023; Heidrich et al., 2020; Stein et al., 2014). Plant diversity benefits from stand-scale microclimatic heterogeneity (Deák et al., 2021; Helbach et al., 2022), and the majority of plant species occurring in central European forests are not adapted to deep shade, but semi-open conditions (Czyżewski and Svenning, 2025). Even forest plant species that can outlive deep shade conditions benefit from microclimatic changes caused by stand-scale disturbances (Messier et al. 2009; von Oheimb and Härdtle 2009).

Ectotherm animals rely on keeping their operational temperature within a range between delayed physiological processes and heat stress (Colinet et al., 2015; Huey et al., 2012). Heterogeneous thermal environments can facilitate thermoregulatory behavior, i.e., moving to optimal thermal microenvironments, of arthropods (Pincebourde and Suppo, 2016; Sears et al., 2011). Biodiversity and performance of arthropod taxa can both be negatively impacted by heat stress (Asch et al., 2025; Junggebauer et al., 2024), or higher in warmer than in colder environments (Lettenmaier et al., 2022; Rothacher et al., 2025; Woods et al., 2015), depending on the specific taxonomic group. For several arthropod groups, there is evidence for thermal heterogeneity on different spatial scales positively affecting their biodiversity and performance (Massó Estaje et al., 2025; Seibold et al., 2016; Terlau et al., 2023). Thus, it is plausible that the heterogeneous microclimatic conditions are one factor driving forest structure-biodiversity relationships of both plants and arthropods.

However, other mechanisms might also come into play. In our design, average microclimate *per se* and its heterogeneity are correlated, making their effects impossible to distinguish (Dormann et al., 2020). In large canopy gaps, not only temperature and its small-scale heterogeneity, but also many other abiotic and biotic factors differ from those in closed-canopy forests. The most obvious example is light availability instead of temperature as a driver of plant diversity (Degen et al., 2005; Helbach et al., 2022). Experimental evidence separating the naturally correlated factors light and temperature implies that light, not temperature, drives plant community responses to canopy openings (De Pauw et al., 2022; Xu et al., 2023). This makes it more likely that, also in our experiment, vascular plant diversity was responding to the within-plot heterogeneity of light, not temperature. Furthermore, altered mineralization rates and nutrient availability in the soil after the interventions (Kovács et al., 2018; Scharenbroch and Bockheim, 2007; Schwarz et al., 2025) could be another contributing factor to the higher vascular plant diversity in gaps. Since many species-rich groups within Diptera and Hymenoptera, e.g., pollinators like bees and hoverflies, rely directly on plants, and Dipteran and Hymenopteran diversity often increase with plant diversity (Scherber et al., 2014; Sobek et al., 2009), it is plausible that higher trophic levels (arthropods) followed the change at the lower trophic level (plants) as a bottom-up diversity effect in gaps (Scherber et al., 2010; Schuldt et al., 2019; Wang et al., 2025). Finally, the link between microclimate and understory plant diversity is not necessarily a one-directional causal relationship, but can be a mutual feedback mechanism, with vascular plant diversity and microclimate influencing each other (Beugnon et al., 2024; Huang et al., 2024; Schnabel et al., 2025; Steinparzer et al., 2025; Wright and Francia, 2024). These complex interacting mechanisms are impossible to disentangle within this framework and will need further investigation. Nevertheless, our results suggest that small-scale microclimatic heterogeneity could be an important contributing factor to the increased biodiversity often observed in canopy openings (Degen et al., 2005; Perlík et al., 2023; Scherber et al., 2014; Shevchenko et al., 2021).

Microclimatic heterogeneity in the tree layer, both vertical and horizontal, was not associated with biodiversity in this study. This is likely because we only sampled arthropods close to the ground, using pitfall traps for spiders and Malaise traps for Diptera and Hymenoptera. However, communities of these groups are often highly vertically stratified in forests (Aikens and Buddle, 2012; Birtele and Hardersen, 2012; Oguri et al., 2014; Sobek et al., 2009). Future studies should combine measuring microclimate and biodiversity in higher layers of forests to test whether microclimatic heterogeneity-diversity relationships also hold in the canopy (Hagge et al., 2024; Wildermuth et al., 2023).

### 4.4 Conclusions

In conclusion, our results demonstrate that standardized point measurements only poorly represent the microclimatic variability within forest plots, but forest structure can be used to predict vertical and horizontal microclimatic heterogeneity. With canopy gaps providing a highly heterogeneous mosaic of warmer and cooler microhabitats in close proximity at the forest floor, and featuring increased biodiversity of vascular plants, Diptera, and Hymenoptera, our results challenge the paradigm of maximized temperature buffering through management towards dense canopies as the only approach to conserve forest biodiversity during climate change. However, with increasing structural complexity and canopy cover, vertical microclimatic heterogeneity increased, potentially affecting the vertical stratification of communities and providing deep shade and cool microhabitats at the forest floor. Resulting from this trade-off between horizontal and vertical microclimatic heterogeneity, forest management approaches that provide both dense, structurally complex stands, and canopy gaps, are most likely to harbor a high diversity of microclimatic habitats, and hence, enable the co- existence of forest organisms with different microclimatic requirements.

## 5 Acknowledgements

We would like to express our sincere gratitude to the Deutsche Forschungsgemeinschaft for funding SP1 (459717468; 459717468; 459717468) within the BETA-FOR Research Unit (FOR 5220/0). JR acknowledges additional funding from the Bavarian State Ministry for Food, Agriculture, Forestry and Tourism (StMELF) (grant no. L062). NE and SC acknowledges support of the German Centre for Integrative Biodiversity Research (iDiv) Halle-Jena-Leipzig iDiv funded by the German Research Foundation (DFG– FZT 118, 202548816) and funding by the DFG (459717468). Furthermore, we would like to thank Hans Stark and the University Forest Sailershausen, as well as the National Park Bavarian Forest, for allowing our experiment in their forests and supporting us tremendously in many ways. We are grateful for the beautiful illustration by Rabea Klümpers, the soil assessment by Hans-Jürgen Gulder, the soil texture analysis by Birgit Terhorst and Anke Pickert, and the amazing hard work in the field and lab of Pierre-André Waite, Felix Bersch, Heidi Inderwies, Nahja Busse, Dimitry Savransky, Biyun Wu, Stefano Hofman-Röckelein, and Joachim Degenfelder.

## 6 Author contributions

**Conceptualization:** Kerstin Pierick, Dominik Seidel, Christian Ammer, Bernhard Schuldt **Methodology:** Kerstin Pierick, Dominik Seidel, Orsi Decker, Martin Ehbrecht, Roman M. Link, Julia Rothacher, Jörg Müller, Michael Köhler, Bernhard Schuldt

**Formal analysis**: Kerstin Pierick, Roman M. Link, Julia Rothacher

**Investigation:** Kerstin Pierick, Dominik Seidel, Orsi Decker, Roman M. Link, Julia Rothacher, Clara Wild, Jean-Léonard Stör, Michael Junginger, Pia Bradler, Sebastian Dittrich, Ludwig Lettenmaier, Lena Rugen, Guido Scholz, Soumen Mallick, Michael Köhler

**Writing – original draft:** Kerstin Pierick

**Writing – review and editing**: Kerstin Pierick, Dominik Seidel, Orsi Decker, Martin Ehbrecht, Roman M. Link, Julia Rothacher, Clara Wild, Jean-Léonard Stör, Michael Junginger, Jörg Müller, Pia Bradler, Benjamin Delory, Nico Eisenhauer, Andreas Fichtner, Sebastian Dittrich, Kim K. Weißing, Ludwig Lettenmaier, Lena Rugen, Guido Scholz, Soumen Mallick, Michael Köhler, Simone Cesarz, Goddert von Oheimb, Christan Ammer, Bernhard Schuldt

**Visualization:** Kerstin Pierick, Kim K. Weißing

**Supervision**: Dominik Seidel, Jörg Müller, Andreas Fichtner, Simone Cesarz, Goddert von Oheimb, Christian Ammer, Bernhard Schuldt

**Funding acquisition**: Dominik Seidel, Jörg Müller, Andreas Fichtner, Simone Cesarz, Nico Eisenhauer, Goddert von Oheimb, Christian Ammer, Bernhard Schuldt

## 7 Data and code availability

All data and R scripts will be made publicly available at the Zenodo community of the BETA- FOR research unit (https://zenodo.org/communities/beta-for/) and will receive a DOI upon acceptance of the article.

