## Supplement for "Microclimatic heterogeneity is associated with forest structural complexity and biodiversity"

#### Figures

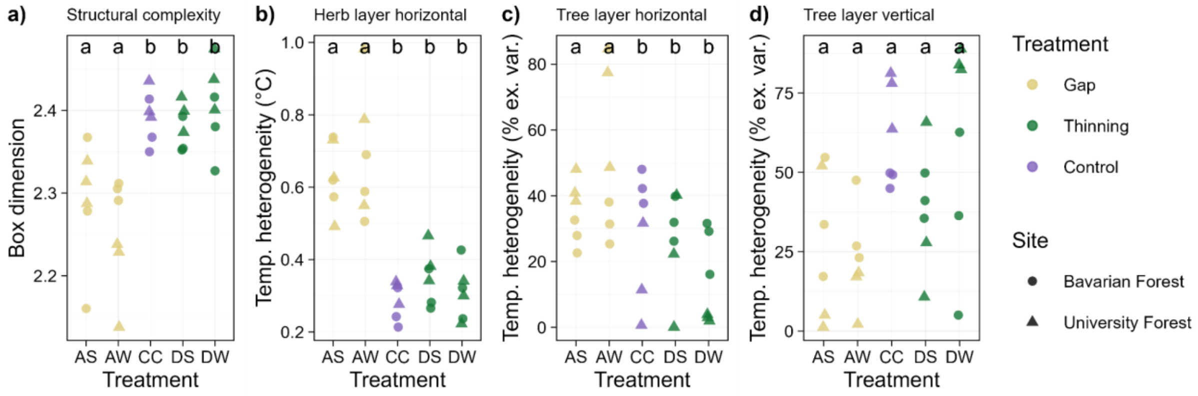


Figure S 1: Effect of non-pooled treatments on key response variables. As opposed to all other results, here, the treatments Gap + Deadwood (AS), Gap (AW), Control (CC), Thinning + Deadwood (DS) and Thinning (DW) are not summarized into Gap, Control and Thinning. The results from the post-hoc Dunn's tests (after Kruskal-Wallis tests), with significant differences between groups indicated by the letters, show that the presence or absence of standing deadwood did not significantly affect the response variables.

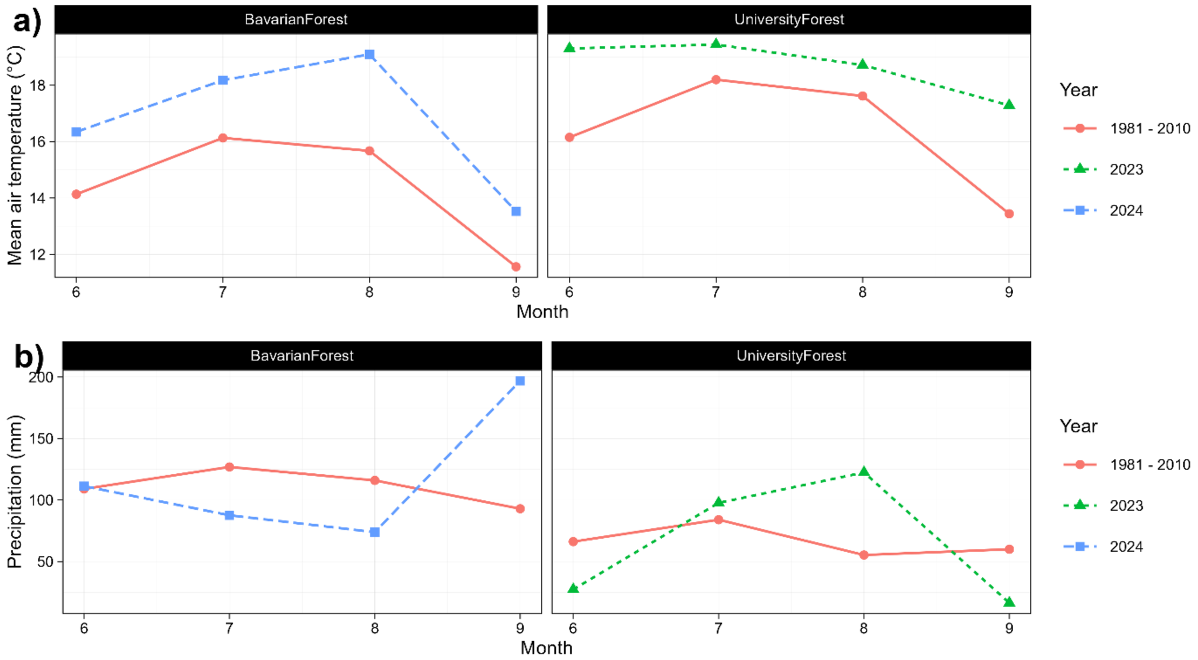


Figure S 2: Weather during the measuring periods and long-term climate at the study sites. We extracted the data for the locations of all forest plots and calculated the means for all plots from grids provided by Deutscher Wetterdienst (DWD Climate Data Center (CDC), 2025a, 2025b, 2025c, 2025d)).

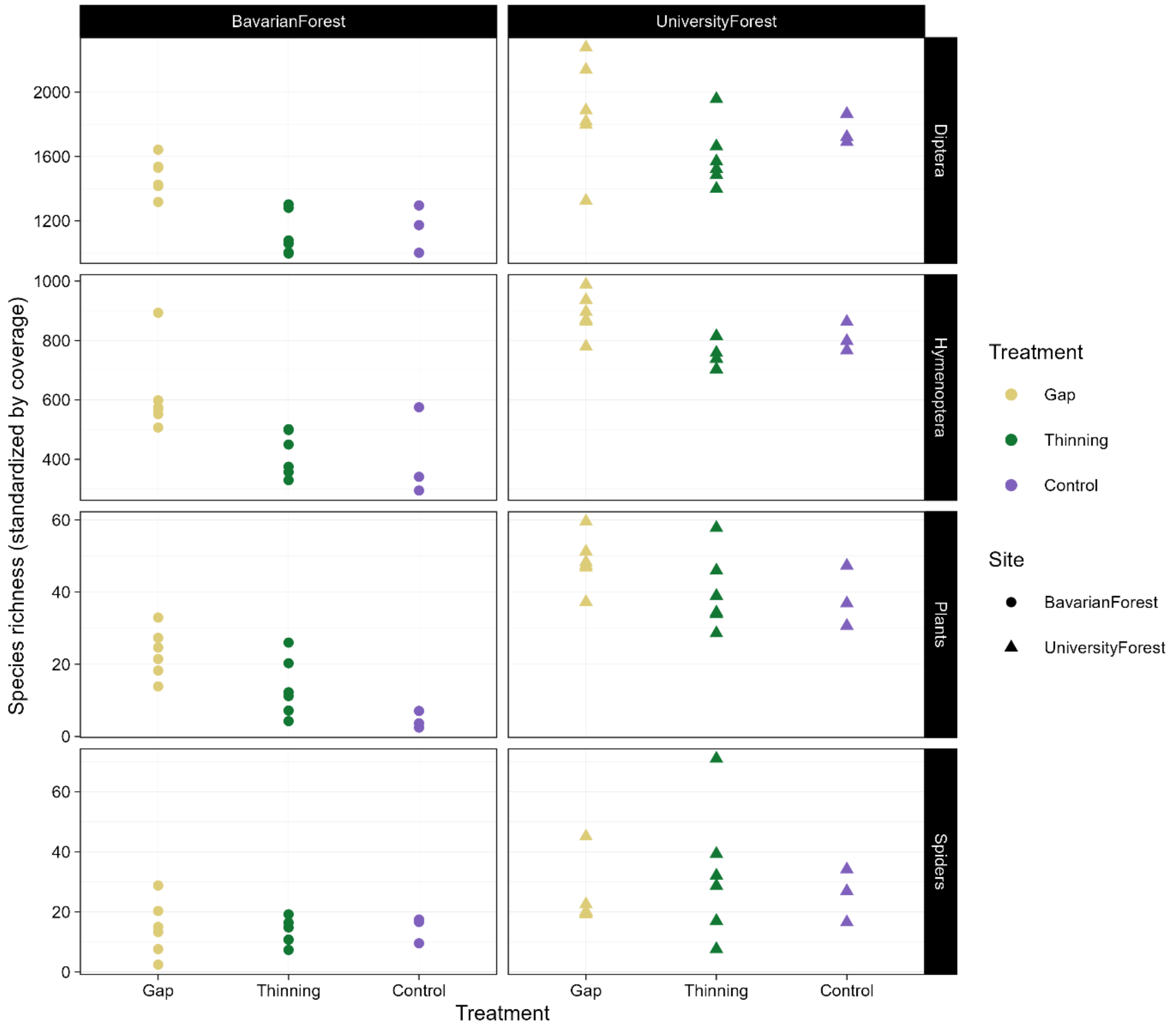


Figure S 3: Biodiversity data before site-correction.

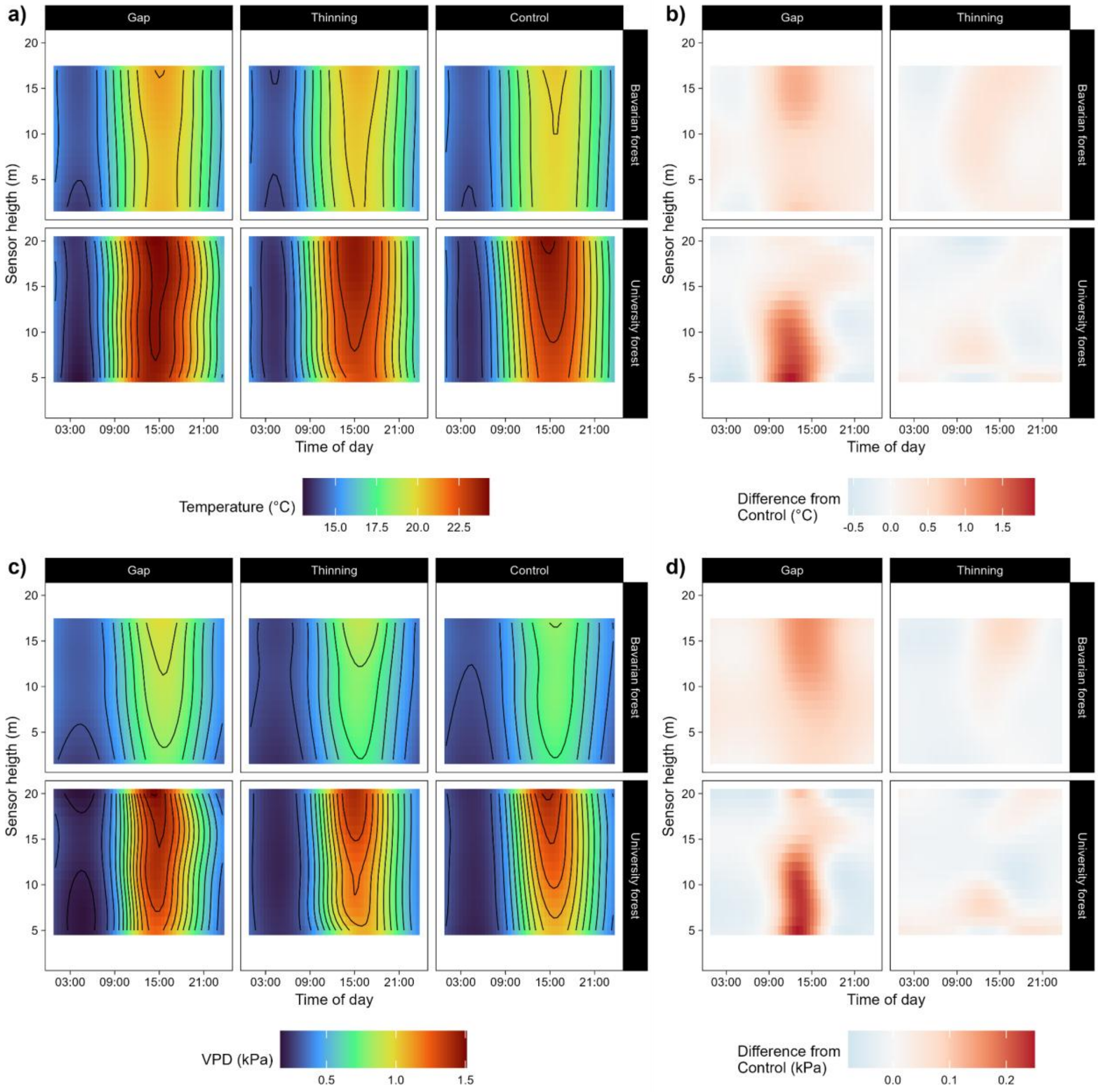


Figure S 4: Average air temperature and vapor pressure deficit (VPD), predicted for different heights and treatments throughout an average summer day. The predictions stem from generalized additive models (GAM). a) shows the absolute predictions, b) shows how the predictions for the Gap and Thinning treatments differ from the Control treatment.

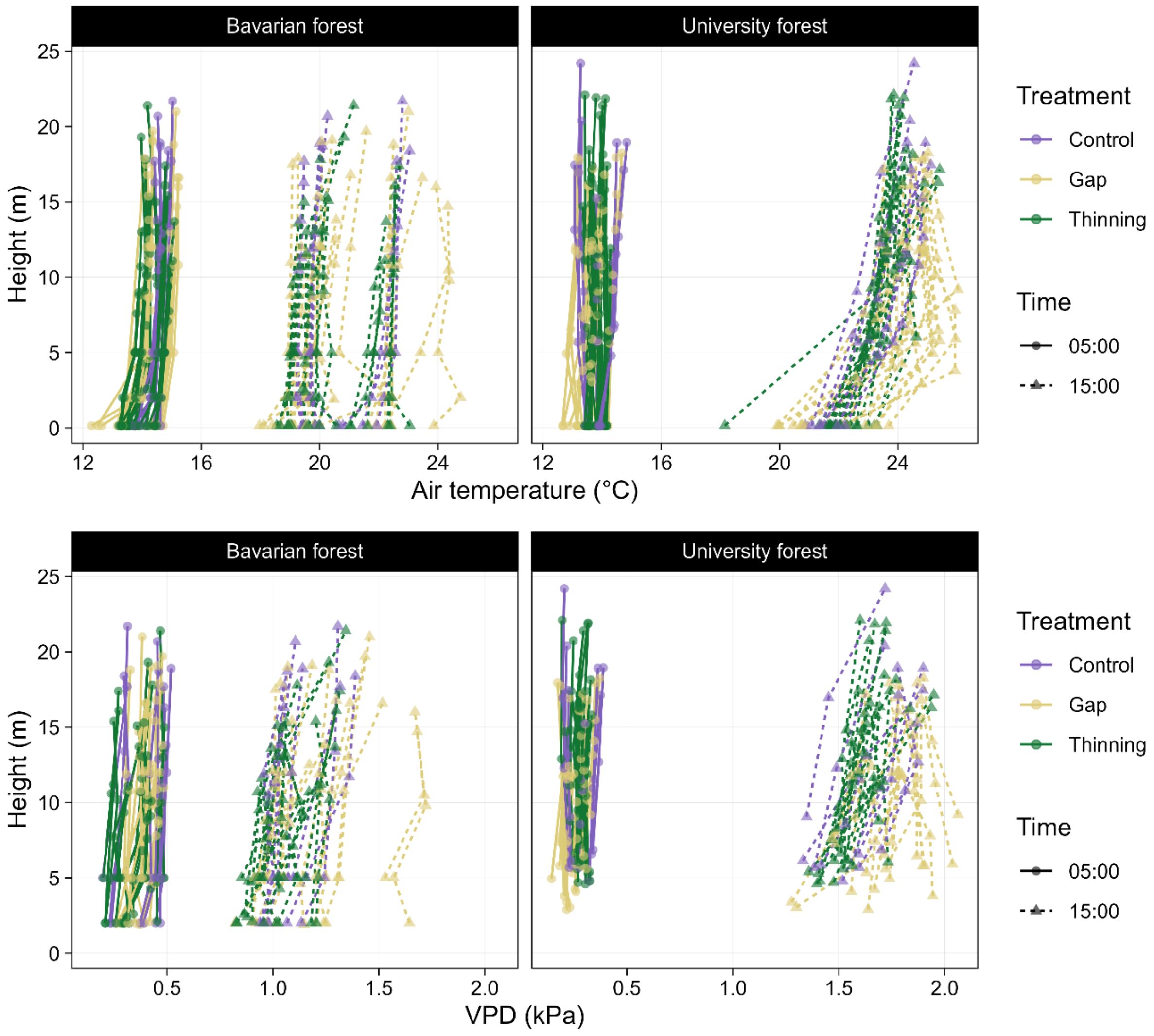


Figure S 5: All individual vertical gradients of air temperature and vapor pressure deficit (VPD). The data points connected by each line represent the Hobo loggers installed on one rope. Shown are the mean VPD values at 05:00 CET and 15:00 CET, averaged over the whole measuring period.

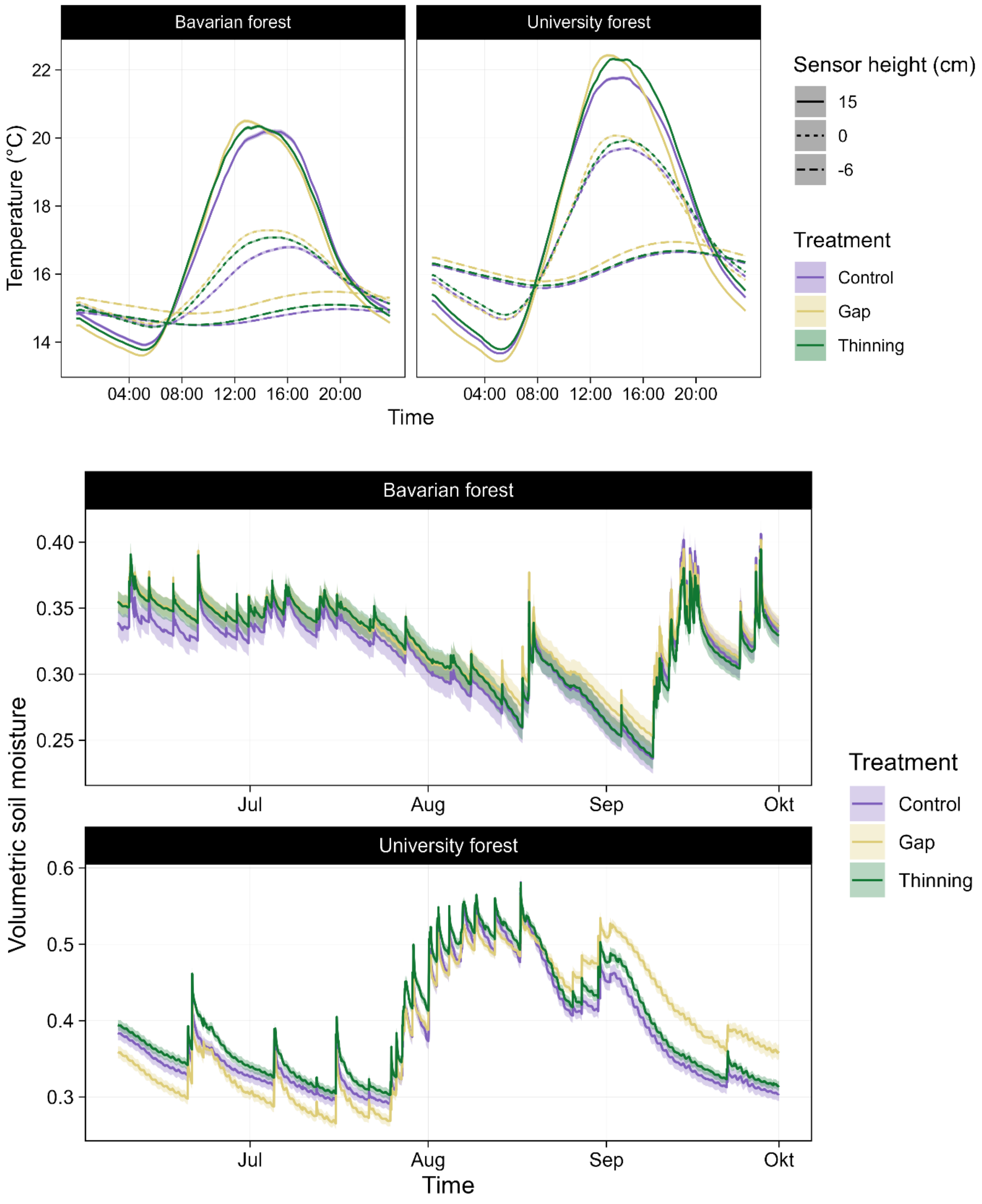


Figure S 6: Summarized data from the Tomst loggers. a) Diurnal temperature oscillations close to the soil surface in different heights, sites, and treatments. The data were measured with Tomst TMS-4 loggers and averaged over all plots, loggers, and days of the measurement periods. The standard error of the mean is indicated by ribbons, which are so narrow they are barely visible due to the high sample size. b) Average volumetric soil moisture throughout the measurement periods (University Forest: 2023, Bavarian Forest: 2024). The data were averaged over all plots and loggers. Ribbons indicate the standard error of the mean.

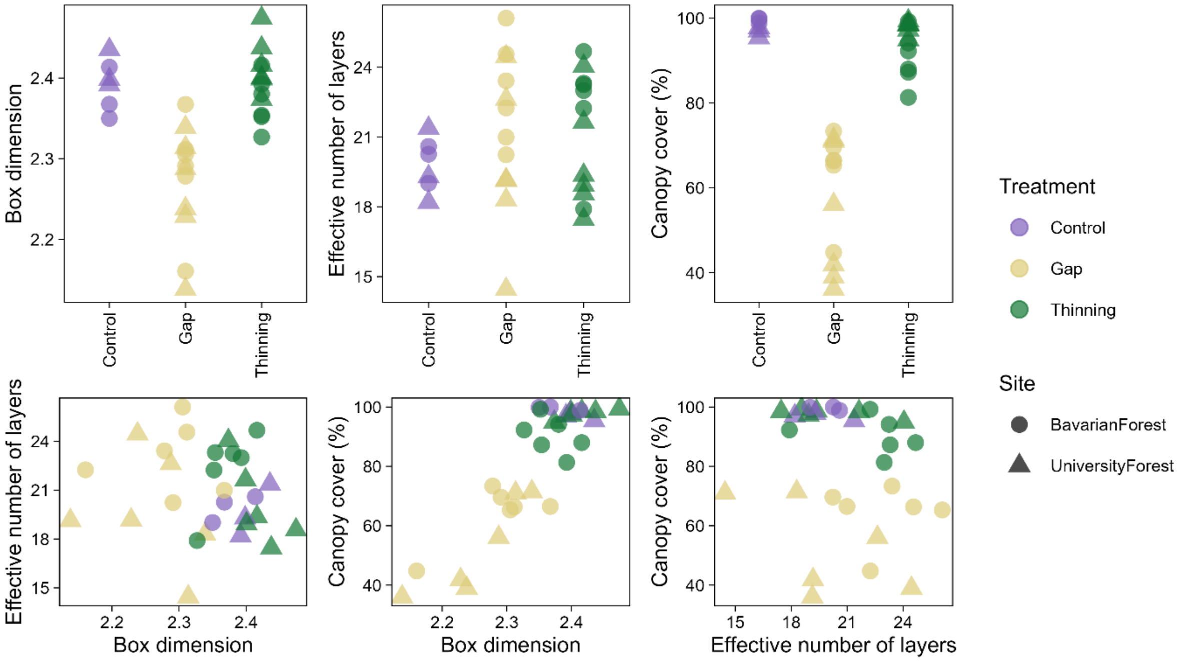


Figure S7: Forest structure indices by treatment, and bivariate relationships between indices.

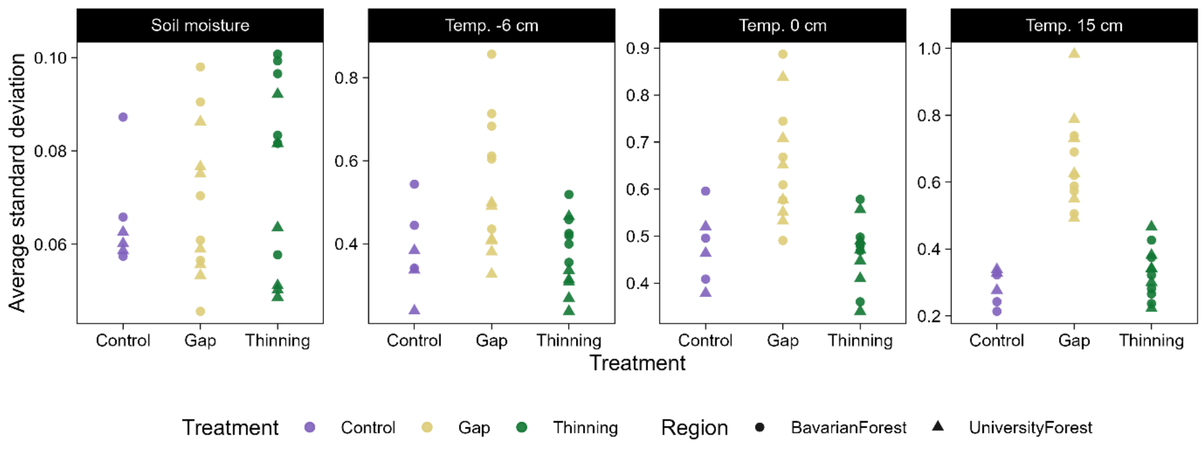


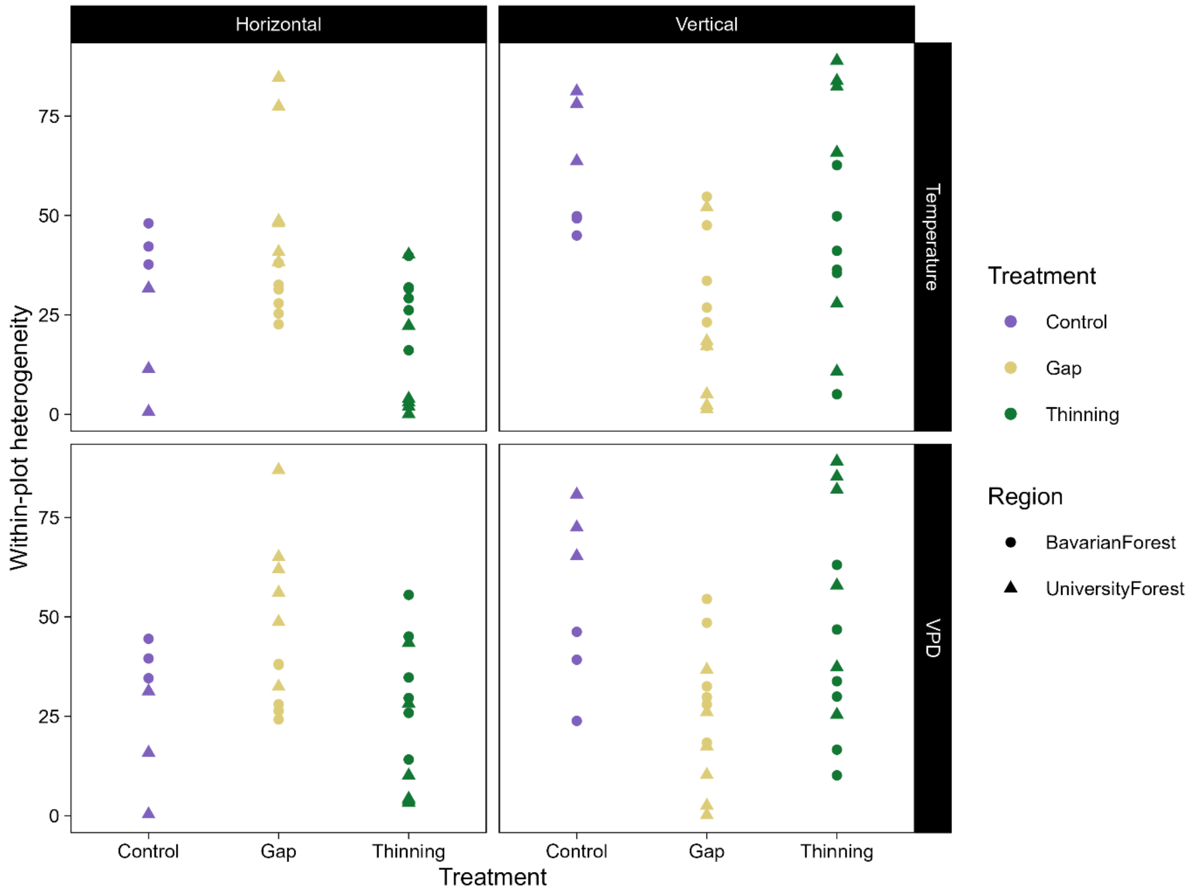


Figure S8: Microclimatic heterogeneity in dependency of treatment. a) Horizontal microclimatic heterogeneity close to the soil surface by treatment. The data were measured with Tomst TMS-4 loggers, and heterogeneity was quantified as the average standard deviation of simultaneously measured values. b) Horizontal and vertical microclimatic heterogeneity of air temperature and vapor pressure deficit (VPD) by treatment. The data were measured with Onset Hobo Mx loggers on ropes, and heterogeneity was quantified with a variance decomposition approach.

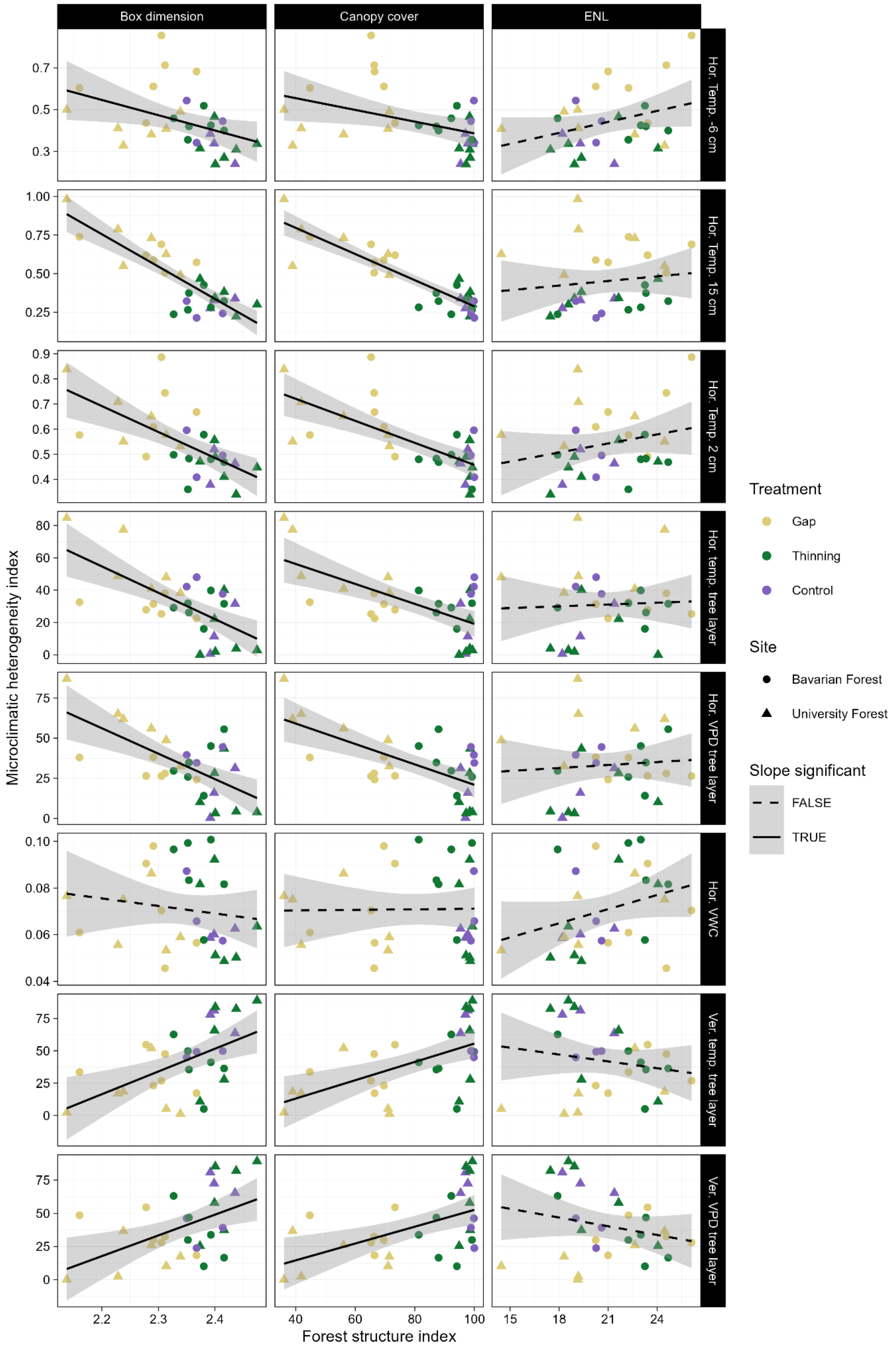


Figure S 9: Relationships between forest structure indices and microclimatic heterogeneity indices. The lines and ribbons show predictions and 95 % confidence intervals from linear models.

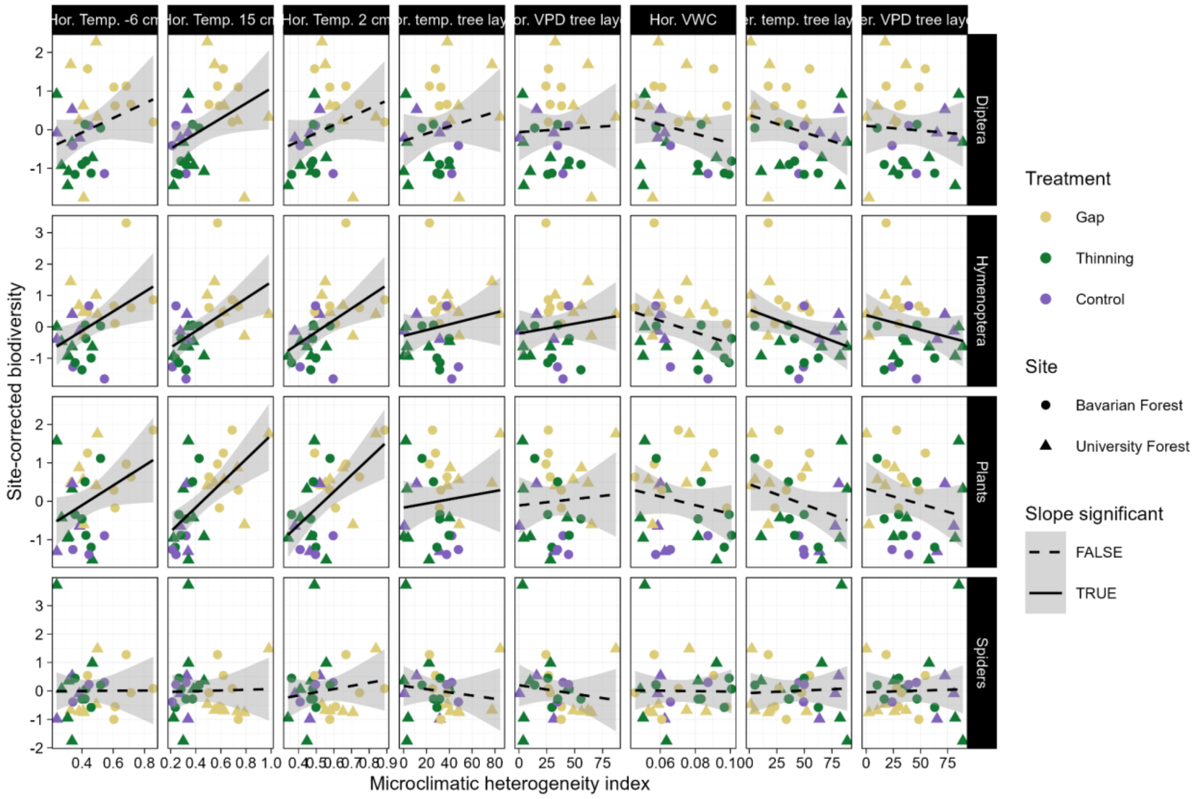


Figure S10: Relationships between microclimatic heterogeneity indices and biodiversity of four taxonomic groups. The lines and ribbons show predictions and 95 % confidence intervals from linear models.

### Tables

Table S 1: Effects of treatment on forest structure indices and microclimatic heterogeneity. X2, df and p are results from Kruskal Wallis tests. Contrast, Estimate and p are results from post-hoc Dunn tests. Significant differences are highlighted in bold font.

| Model | $X^2$ | df | p | Contrast | Estimate | p |
|---|---|---|---|---|---|---|
| | | | *Forest structure indices* | | | |
| **Box dimension** | 19.03 | 2 | **< 0.0001** | **Control-Gap** | 3.13 | **0.0008** |
| | | | | Control-Thinning | -0.13 | 0.4 |
| | | | | **Gap-Thinning** | -4.03 | **< 0.0001** |
| Eff. number of layers | 1.65 | 2 | 0.4 | | | |
| **Canopy cover** | 21.84 | 2 | **< 0.0001** | **Control-Gap** | 4.05 | **< 0.0001** |
| | | | | Control-Thinning | 0.97 | 0.2 |
| | | | | **Gap-Thinning** | -3.78 | **0.0001** |
| | | | *Tomst logger-derived microclimatic heterogeneity* | | | |
| **Hor. temp. het. -6 cm** | 7.14 | 2 | **0.03** | **Control-Gap** | -1.87 | **0.03** |
| | | | | Control-Thinning | 0.17 | 0.4 |
| | | | | **Gap-Thinning** | 2.50 | **0.006** |
| **Hor. temp. het. 0 cm** | 15.24 | 2 | **0.0005** | **Control-Gap** | -2.73 | **0.003** |
| | | | | Control-Thinning | 0.27 | 0.4 |
| | | | | **Gap-Thinning** | 3.66 | **0.0001** |
| **Hor. temp. het. 15 cm** | 21.45 | 2 | **< 0.0001** | **Control-Gap** | -3.90 | **< 0.0001** |
| | | | | Control-Thinning | -0.74 | 0.2 |
| | | | | **Gap-Thinning** | 3.87 | **0.0001** |
| Hor. VWC het. | 0.88 | 2 | 0.7 | | | |
| | | | *Hobo logger-derived microclimatic heterogeneity* | | | |
| **Hor. temp. tree layer** | 6.87 | 2 | **0.03** | Control-Gap | -1.02 | 0.2 |
| | | | | Control-Thinning | 1.12 | 0.1 |
| | | | | **Gap-Thinning** | 2.62 | **0.004** |
| **Ver. temp. tree layer** | 8.84 | 2 | **0.01** | **Control-Gap** | 2.71 | **0.003** |
| | | | | Control-Thinning | 0.89 | 0.2 |

| | | | | | | |
|---|---|---|---|---|---|---|
| | | | | **Gap-Thinning** | -2.23 | **0.01** |
| Hor. VPD tree layer | 4.48 | 2 | 0.1 | | | |
| **Ver. VPD tree layer** | 6.78 | 2 | **0.03** | **Control-Gap** | 2.27 | **0.01** |
| | | | | Control-Thinning | 0.57 | 0.3 |
| | | | | **Gap-Thinning** | -2.09 | **0.02** |

Table S 2: Effects forest structure indices on microclimatic heterogeneity indices. Shown are results from linear models. Significant differences are highlighted in bold font.

| Response | Predictor | $R^2$ | Parameter | Estimate | p |
|---|---|---|---|---|---|
| **Hor. temp. het. -6 cm** | **Box dimension** | 0.16 | Intercept | 2.16 | |
| | | | Slope | -0.73 | **0.03** |
| | | | Res. std. error | 0.13 | |
| **Hor. temp. het. -6 cm** | **Canopy cover** | 0.17 | Intercept | 0.67 | |
| | | | Slope | -0.003 | **0.02** |
| | | | Res. std. error | 0.14 | |
| Hor. temp. het. -6 cm | ENL | 0.11 | Intercept | 0.07 | |
| | | | Slope | 0.02 | 0.08 |
| | | | Res. std. error | 0.14 | |
| **Hor. temp. het. 0 cm** | **Box dimension** | 0.39 | Intercept | 2.96 | |
| | | | Slope | -1.03 | **0.0002** |
| | | | Res. std. error | 0.10 | |
| **Hor. temp. het. 0 cm** | **Canopy cover** | 0.49 | Intercept | 0.90 | |
| | | | Slope | -0.004 | **<0.0001** |
| | | | Res. std. error | 0.09 | |
| Hor. temp. het. 0 cm | ENL | 0.06 | Intercept | 0.29 | |
| | | | Slope | 0.01 | 0.2 |
| | | | Res. std. error | 0.13 | |
| **Hor. temp. het. 15 cm** | **Box dimension** | 0.70 | Intercept | 5.37 | |
| | | | Slope | -2.20 | **<0.0001** |
| | | | Res. std. error | 0.11 | |
| **Hor. temp. het. 15 cm** | **Canopy cover** | 0.79 | Intercept | 1.14 | |
| | | | Slope | -0.008 | **<0.0001** |
| | | | Res. std. error | 0.09 | |
| Hor. temp. het. 15 cm | ENL | 0.02 | Intercept | 0.24 | |
| | | | Slope | 0.01 | 0.5 |
| | | | Res. std. error | 0.20 | |
| Hor. VWC het. | Box dimension | 0.02 | Intercept | 0.15 | |
| | | | Slope | -0.03 | 0.4 |
| | | | Res. std. error | 0.12 | |
| Hor. VWC het. | Canopy cover | <0.01 | Intercept | 0.01 | |
| | | | Slope | <0.001 | 0.9 |
| | | | Res. std. error | 0.02 | |
| Hor. VWC het. | ENL | 0.01 | Intercept | 0.01 | |
| | | | Slope | 0.002 | 0.09 |
| | | | Res. std. error | 0.02 | |
| **Hor. temp. tree layer** | **Box dimension** | 0.42 | Intercept | 423.5 | |
| | | | Slope | -163.9 | **0.0001** |
| | | | Res. std. error | 15.5 | |
| **Hor. temp. tree layer** | **Canopy cover** | 0.41 | Intercept | 80.9 | |
| | | | Slope | -0.62 | **0.0001** |
| | | | Res. std. error | 15.5 | |
| Hor. temp. tree layer | ENL | <0.01 | Intercept | 23.4 | |
| | | | Slope | 0.37 | 0.8 |
| | | | Res. std. error | 20.3 | |
| **Ver. temp. tree layer** | **Box dimension** | 0.28 | Intercept | -370.7 | |
| | | | Slope | 176.0 | **0.003** |
| | | | Res. std. error | 22.7 | |
| **Ver. temp. tree layer** | **Canopy cover** | 0.31 | Intercept | -15.3 | |
| | | | Slope | 0.71 | **0.001** |
| | | | Res. std. error | 22.2 | |
| Ver. temp. tree layer | ENL | 0.03 | Intercept | 78.9 | |
| | | | Slope | -1.77 | 0.3 |
| | | | Res. std. error | 26.3 | |

| | | | | | |
|---|---|---|---|---|---|
| **Hor. VPD tree layer** | **Box dimension** | 0.39 | Intercept | 405.2 | |
| | | | Slope | -158.7 | **0.0002** |
| | | | Res. std. error | 15.9 | |
| **Hor. VPD tree layer** | **Canopy cover** | 0.44 | Intercept | 84.6 | |
| | | | Slope | -0.64 | **<0.0001** |
| | | | Res. std. error | 15.3 | |
| Hor. VPD tree layer | ENL | 0.01 | Intercept | 20.3 | |
| | | | Slope | 0.62 | 0.7 |
| | | | Res. std. error | 20.3 | |
| **Ver. VPD tree layer** | **Box dimension** | 0.24 | Intercept | -325.5 | |
| | | | Slope | 156.0 | **0.006** |
| | | | Res. std. error | 22.2 | |
| **Ver. VPD tree layer** | **Canopy cover** | 0.28 | Intercept | -10.7 | |
| | | | Slope | 0.63 | **0.003** |
| | | | Res. std. error | 21.7 | |
| Ver. VPD tree layer | ENL | 0.05 | Intercept | 86.2 | |
| | | | Slope | -2.19 | 0.2 |
| | | | Res. std. error | 24.8 | |

Table S 3: Effects microclimatic heterogeneity indices on biodiversity of plants, spiders, Diptera and Hymenoptera. Shown are results from linear models. Significant differences are highlighted in bold font.

| Response | Predictor | $R^2$ | Parameter | Estimate | p |
|---|---|---|---|---|---|
| **Plants** | **Hor. temp. het. -6 cm** | 0.14 | Intercept | -1.14 | |
| | | | Slope | 2.58 | **0.04** |
| | | | Res. std. error | 0.95 | |
| **Plants** | **Hor. temp. het. 0 cm** | 0.32 | Intercept | -2.35 | |
| | | | Slope | 4.34 | **0.001** |
| | | | Res. std. error | 0.84 | |
| **Plants** | **Hor. temp. het. 15 cm** | 0.39 | Intercept | -1.40 | |
| | | | Slope | 3.14 | **0.0003** |
| | | | Res. std. error | 0.80 | |
| Plants | Hor. VWC het. | 0.04 | Intercept | 0.80 | |
| | | | Slope | -11.2 | 0.3 |
| | | | Res. std. error | 0.10 | |
| Plants | Hor. temp. tree layer | 0.01 | Intercept | -0.17 | |
| | | | Slope | 0.01 | 0.6 |
| | | | Res. std. error | 1.01 | |
| Plants | Ver. temp. tree layer | 0.08 | Intercept | 0.44 | |
| | | | Slope | -0.01 | 0.1 |
| | | | Res. std. error | 0.98 | |
| Plants | Hor. VPD tree layer | <0.01 | Intercept | -0.11 | |
| | | | Slope | 0.003 | 0.7 |
| | | | Res. std. error | 1.02 | |
| Plants | Ver. VPD tree layer | 0.04 | Intercept | 0.32 | |
| | | | Slope | -0.01 | 0.3 |
| | | | Res. std. error | 1.00 | |
| Spiders | Hor. temp. het. -6 cm | 0.04 | Intercept | -0.02 | |
| | | | Slope | 0.04 | 0.9 |
| | | | Res. std. error | 1.02 | |
| Spiders | Hor. temp. het. 0 cm | 0.02 | Intercept | -0.61 | |
| | | | Slope | 1.12 | 0.4 |
| | | | Res. std. error | 1.01 | |
| Spiders | Hor. temp. het. 15 cm | <0.01 | Intercept | -0.06 | |
| | | | Slope | 0.12 | 0.9 |
| | | | Res. std. error | 1.02 | |
| Spiders | Hor. VWC het. | <0.01 | Intercept | 0.06 | |
| | | | Slope | -0.79 | 0.9 |
| | | | Res. std. error | 1.02 | |
| Spiders | Hor. temp. tree layer | 0.01 | Intercept | 0.18 | |
| | | | Slope | -0.01 | 0.6 |
| | | | Res. std. error | 1.01 | |
| Spiders | Ver. temp. tree layer | <0.01 | Intercept | -0.07 | |
| | | | Slope | 0.001 | 0.8 |

| | | | | | |
|---|---|---|---|---|---|
| | | | Res. std. error | 1.02 | |
| Spiders | Hor. VPD tree layer | 0.01 | Intercept | 0.20 | |
| | | | Slope | -0.01 | 0.5 |
| | | | Res. std. error | 1.01 | |
| Spiders | Ver. VPD tree layer | <0.01 | Intercept | -0.05 | |
| | | | Slope | 0.001 | 0.9 |
| | | | Res. std. error | 1.02 | |
| Diptera | Hor. temp. het. -6 cm | 0.07 | Intercept | -0.83 | |
| | | | Slope | 1.88 | 0.1 |
| | | | Res. std. error | 0.98 | |
| Diptera | Hor. temp. het. 0 cm | 0.08 | Intercept | -1.14 | |
| | | | Slope | 2.11 | 0.1 |
| | | | Res. std. error | 0.98 | |
| **Diptera** | **Hor. temp. het. 15 cm** | 0.15 | Intercept | -0.88 | |
| | | | Slope | 1.95 | **0.04** |
| | | | Res. std. error | 0.94 | |
| Diptera | Hor. VWC het. | 0.04 | Intercept | 0.84 | |
| | | | Slope | -11.9 | 0.3 |
| | | | Res. std. error | 1.00 | |
| Diptera | Hor. temp. tree layer | 0.01 | Intercept | -0.29 | |
| | | | Slope | 0.01 | 0.3 |
| | | | Res. std. error | 1.00 | |
| Diptera | Ver. temp. tree layer | 0.06 | Intercept | 0.38 | |
| | | | Slope | -0.01 | 0.2 |
| | | | Res. std. error | 0.99 | |
| Diptera | Hor. VPD tree layer | <0.01 | Intercept | -0.07 | |
| | | | Slope | 0.002 | 0.8 |
| | | | Res. std. error | 1.02 | |
| Diptera | Ver. VPD tree layer | <0.01 | Intercept | 0.10 | |
| | | | Slope | -0.002 | 0.7 |
| | | | Res. std. error | 1.02 | |
| Hymenoptera | Hor. temp. het. -6 cm | 0.20 | Intercept | -1.36 | |
| | | | Slope | 3.09 | 0.01 |
| | | | Res. std. error | 0.91 | |
| **Hymenoptera** | **Hor. temp. het. 0 cm** | 0.24 | Intercept | -2.03 | |
| | | | Slope | 3.73 | **0.006** |
| | | | Res. std. error | 0.89 | |
| **Hymenoptera** | **Hor. temp. het. 15 cm** | 0.27 | Intercept | 1.18 | |
| | | | Slope | 2.61 | **0.004** |
| | | | Res. std. error | 0.87 | |
| Hymenoptera | Hor. VWC het. | 0.10 | Intercept | 1.29 | |
| | | | Slope | -18.2 | 0.09 |
| | | | Res. std. error | 0.97 | |
| Hymenoptera | Hor. temp. tree layer | 0.03 | Intercept | -0.28 | |
| | | | Slope | 0.01 | 0.3 |
| | | | Res. std. error | 1.00 | |
| Hymenoptera | Ver. temp. tree layer | 0.12 | Intercept | 0.55 | |
| | | | Slope | -0.01 | 0.06 |
| | | | Res. std. error | 0.95 | |
| Hymenoptera | Hor. VPD tree layer | 0.02 | Intercept | -0.21 | |
| | | | Slope | 0.01 | 0.5 |
| | | | Res. std. error | 1.01 | |
| Hymenoptera | Ver. VPD tree layer | 0.05 | Intercept | 0.37 | |
| | | | Slope | -0.01 | 0.2 |
| | | | Res. std. error | 0.99 | |

### Methods

#### Method S1: Soil moisture calibration

We derived volumetric soil water content from the raw output of the loggers using the `myClim::mc_calc_vwc()` function from the `myClim` R package with plot-specific, soil-texture-dependent calibration parameters. For 21 plots, particle size distribution was measured using laser granulometry with a HORIBA Partica LA-960V2 laser diffraction analyzer (HORIBA 2024b). This device applies the Mie scattering principle to measure particles between 0.01 µm and 5000 µm (HORIBA 2024a). Red laser and blue LED light are passed through a suspended sample, and the pattern of scattered light—detected at different angles—is used to calculate particle sizes via mathematical models (HORIBA 2024a). Blue diodes provide higher precision for small particles. Measurement settings for refractive indices followed Özer et al. (2010). Each sample was analyzed twice, giving four measurements in total, and the results were compiled in tables. For the remaining 9 plots (all located at the Bavarian Forest site), this was not possible due to thick organic layers, which is why we used particle size fraction proportions estimated with the soil texture by feel method. Sand, silt and clay proportions of each plot were entered into the calibration tool provided by the manufacturer of the Tomst loggers to acquire soil texture-specific calibration parameters. While the methodological inconsistency in acquiring the calibration parameters hinders meaningful comparisons of soil moisture between plots, the method was consistent within each plot, allowing us to quantify within-plot heterogeneity.

DWD Climate Data Center (CDC), 2025c. Grids of monthly averaged daily air temperature (2m) over Germany v1.0.

DWD Climate Data Center (CDC), 2025d. Grids of monthly total precipitation over Germany v1.0.

Horiba 2024a: A Guidebook to Particle Size Analysis. Irvine, USA.

Horiba 2024b: Partica LA-960V2. Laser Scattering Particle Size Distribution Analyzer. URL: https://www.horiba.com/int/scientific/products/detail/action/show/Product/partica-la-960v2-1944/ (last checked 20.08.2024)

Özer, M., et al. 2010: Effect of Particle Optical Properties on Size Distribution of Soils Obtained by Laser Diffraction. Environmental & Engineering Geoscience 16(2): 163-173.
